# Neural encoding of phrases and sentences in spoken language comprehension

**DOI:** 10.1101/2021.07.09.451747

**Authors:** Fan Bai, Antje S. Meyer, Andrea E. Martin

**Affiliations:** Max Planck Institute for Psycholinguistics, Nijmegen, The Netherlands; Donders Institute for Brain, Cognition, and Behaviour, Radboud University, Nijmegen, The Netherlands

## Abstract

Speech stands out in the natural world as a biological signal that communicates formally-specifiable complex meanings. However, the acoustic and physical dynamics of speech do not injectively mark the linguistic structure and meaning that we perceive. Linguistic structure must therefore be inferred through the human brain’s endogenous mechanisms, which remain poorly understood. Using electroencephalography, we investigated the neural response to synthesized spoken phrases and sentences that were closely physically-matched but differed in syntactic structure, under either linguistic or non-linguistic task conditions. Differences in syntactic structure were well-captured in theta band (∼ 2 to 7 Hz) phase coherence, phase connectivity degree at low frequencies (< ∼ 2 Hz), and in both intensity and degree of power connectivity of induced neural response in the alpha band (∼ 7.5 to 13.5 Hz). Theta-gamma phase-amplitude coupling was found when participants listened to speech, but it did not discriminate between syntactic structures. Spectral-temporal response function modelling suggested different encoding states in both temporal and spectral dimensions as a function of the amount and type of linguistic structure perceived, over and above the acoustically-driven neural response. Our findings provide a comprehensive description of how the brain separates linguistic structures in the dynamics of neural responses, and imply that phase synchronization and strength of connectivity can be used as readouts for constituent structure, providing a novel basis for future neurophysiological research on linguistic structure in the brain.

## Introduction

Speech contains an abundance of acoustic features in both the time and frequency domains (Shannon, Zeng, Kamath, Wygonski, & Ekelid, 1995; Smith, Delgutte, & Oxenham, 2002; Zeng et al., 2005). While these features are crucial for speech comprehension, they do not themselves signpost the linguistic units and structures that give rise to meaning. Spoken language comprehension therefore relies on listeners to go beyond the information given and infer the presence of linguistic structure based on their knowledge of language. As such, many theories posit that linguistic structures – ranging from syllables to ‘words’ to syntactic structures - are constructed via an endogenous inference process (Bever & Poeppel, 2010; Brown, Tanenhaus, & Dilley, 2021; Friederici, 1995; Hagoort, 2013; Halle & Stevens, 1962; Marslen-Wilson & Tyler, 1980; Marslen-Wilson, 1987; Marslen-Wilson & Welsh, 1978; Martin, 2016, 2020; Martin & Doumas, 2017, 2019, 2020; Meyer, Sun, & Martin, 2020; Phillips, 2003; Poeppel & Monahan, 2011). On this view, also known as ‘analysis-by-synthesis’ (Halle, Stevens, Wathen-Dunn, & Woods, 1959), speech triggers internal generation of memory representations (synthesis), which are compared to the sensory input (analysis). This results in linguistic structures that come about through the synthesis of internal brain states (*viz.*, linguistic knowledge) with sensory representations via perceptual inference (Marslen-Wilson & Welsh, 1978; Martin, 2016, 2020). Recent studies have begun to investigate the neural activity that corresponds to the emergence of linguistic structure (e.g., Ding, Melloni, Zhang, Tian, & Poeppel, 2016; Kaufeld et al., 2020; Keitel, Gross, & Kayser, 2018; Meyer & Gumbert, 2018), in particular in terms of temporal and spatial dynamics of brain rhythms. However, despite these efforts, details of how the brain encodes or distinguishes syntactic structures remain unknown. In this study, we investigated the neural responses to minimally different linguistic structures, specifically phrases like ‘*the red vase’* compared to sentences like ‘*the vase is red’*. We investigated which dimensions of neural activity distinguish the linguistic structure of these phrases and sentences, by minimizing their differences in acoustic-energetic/temporal-spectral profiles and semantic components.

### Low frequency neural oscillations and linguistic structure

A growing neuroscience literature suggests that low frequency neural oscillations ( < 8 Hz) are involved in processing linguistic structures (Brennan & Martin, 2020; Brodbeck, Hong, & Simon, 2018; Ding, Melloni, et al., 2017; Ding et al., 2016; Gui et al., 2020; Gwilliams & King, 2020; Jin, Lu, & Ding, 2020; Jin, Zou, Zhou, & Ding, 2018; Kaufeld et al., 2020; Keitel et al., 2018; Martin & Doumas, 2017, 2019; Meyer, 2018; Meyer & Gumbert, 2018; Obleser & Kayser, 2019; Zhou, Melloni, Poeppel, & Ding, 2016). In a highly influential study by Ding and colleagues (2016), low frequency power ‘tagged’ the occurrence of phrases and sentences in controlled speech stimuli. That is, power increases were observed that coincided with occurences of phrases or sentences at a fixed rate of 2 and 1Hz, respectively. Additionally, the grouping of words into phrases of different lengths modulated the location of the frequency tag accordingly, indicating that power at certain frequencies could track linguistic structures. Subsequent research has further confirmed the sensitivity of oscillatory power and phase in the delta band (∼2-4 Hz) to higher level linguistic structures like phrases (Brennan & Martin, 2020; Kaufeld et al., 2020; Meyer & Gumbert, 2018; Meyer, Henry, Gaston, Schmuck, & Friederici, 2017; Peña & Melloni, 2012).

Inspired by these empirical studies, Martin and Doumas (2017) recently provided a theoretical, computationally explicit framework for understanding the role of low frequency neural oscillations in generating linguistic structure. Martin and Doumas (2017) reproduced the frequency tagging results reported by Ding et al. (2016) in an artificial neural network model that uses time (unit firing asynchrony^1^) to encode structural relations between words (see also Martin & Doumas, 2019). Based on their model, Martin and Doumas hypothesized that low frequency power and synchronization should depend on the number of constituents that are represented at a given timestep. In their model’s representational coding scheme, constituents are represented as (localist) relations between distributed representations in time. Thus, the ongoing dynamics of neural ensembles involved in coding linguistic units and their structural relations are what constitute ‘linguistic structure’ in such a neural sytem (Martin, 2016, 2020). The current study tested this prediction using Dutch phrases (e.g., *de rode vas, ‘*the red vase,’ with 2-4 constituents depending on the selection of a formal syntactic denotation; contituents contained determiner, adjectival, and nominal heads) and sentences (e.g., *de vas is rood,* ‘the vase is red,’with 3-10 constituents depending on syntactic account; with determiner, adjectival and nominal heads) that were closely matched in acoustic-physical characteristics but differed in syntactic structure (viz., the number and type of constituents). **Fig 1** shows the syntactic structures in the sample phrase-sentence pair, *de rode vaas (the red vase)* and *de vaas is rood (the vase is red)* in a minimalistic syntactic denotation.

**Fig 1.**
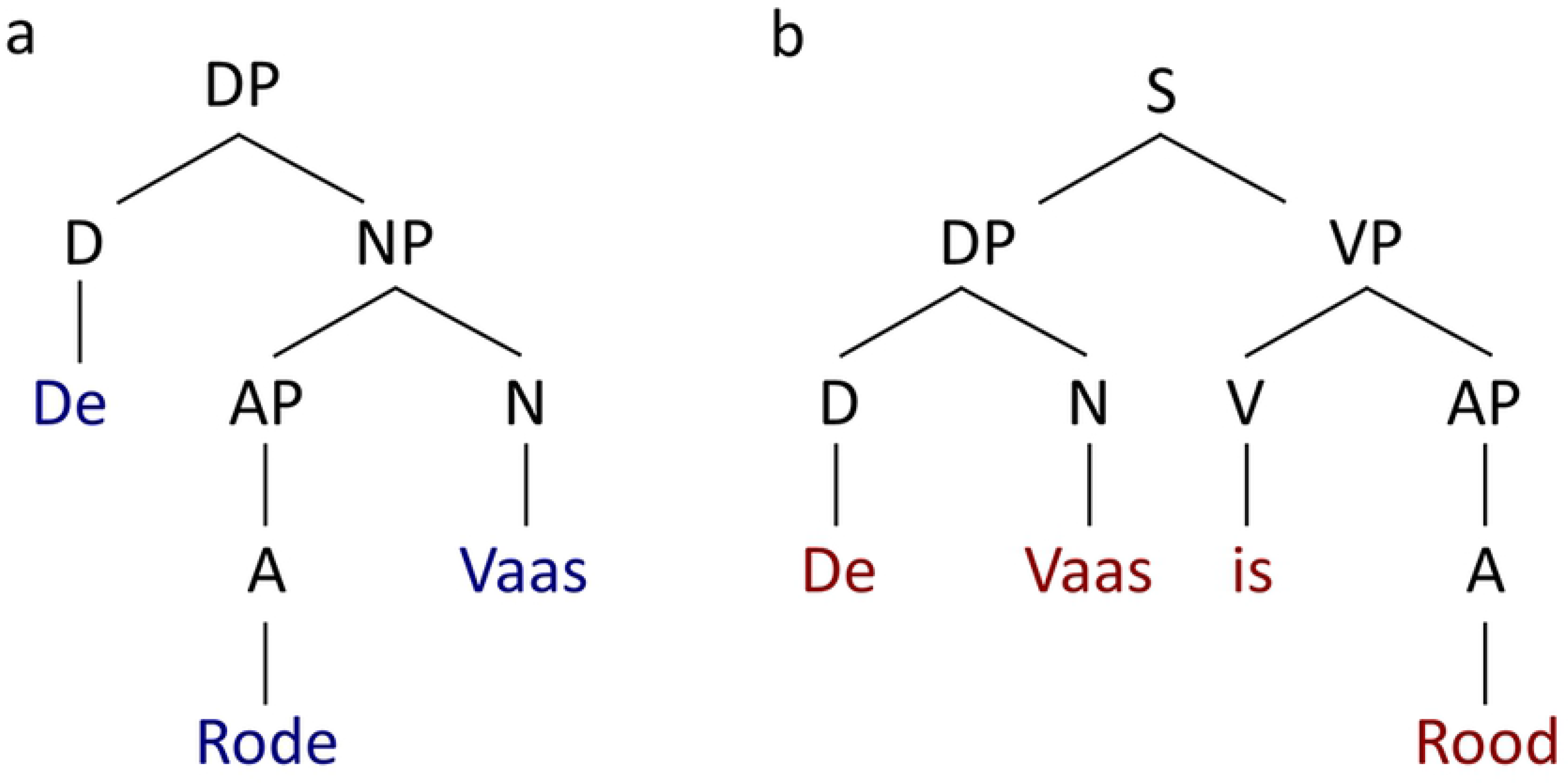
Syntactic structure comparison between phrase and sentence. **(a)** Tree representation of the syntactic structure for the sample phrase, *de rode vaas (the red vase).* The determiner phase (DP) can first be decomposed into a determiner (D) and a noun phrase (NP), then the NP can be separated into an adjective phrase (AP), which equivalents to an adjective (A), and a noun (N). **(b)** Tree representation of the syntactic structure for the sample sentence, *de vaas is rood (the vase is red).* The sentence can be decomposed into two parts, which are a determiner phrase (DP) and a verb phrase (VP), respectively. The DP can then be separateded into a determiner (D) and a noun (N), and the VP can be separated into a combination between a verb (V) and an adjective (A).

### Low frequency oscillations and speech intelligibility

Low frequency neural activity, in particular phase coherence in the ∼2-7 Hz theta band, is highly correlated with spoken language comprehension (e.g., Doelling, Arnal, Ghitza, & Poeppel, 2014; Howard & Poeppel, 2010; Luo & Poeppel, 2007; Peelle, Gross, & Davis, 2013; Zoefel, Archer-Boyd, & Davis, 2018; Zoefel & Van Rullen, 2016). For example, an magnetoencephalography (MEG) study by Luo and Poeppel (2007) showed that theta band phase coherence was positively correlated with speech intelligibility. One possible explanation of such findings is that people’s ability to infer linguistic structure increases with speech intelligibility, with resulting changes in low frequency neural activity (see also Ding et al.).

Low frequency neural oscillations may be especially important for speech processing because they roughly occur at the average syllable rate across various human languages (Ding, Patel, et al., 2017; Pellegrino, Coupé, & Marsico, 2011; Varnet, Ortiz-Barajas, Erra, Gervain, & Lorenzi, 2017). The brain may use syllables, which are abstract linguistic units, as the primitive units to analyze spoken language (Giraud & Poeppel, 2012; Luo & Poeppel, 2007; Poeppel & Assaneo, 2020). Indeed, a view has emerged wherein the brain employs an inherent cortical rhythm at a syllabic rate that can be altered by manipulations of linguistic structure or intelligibility. One possible synthesis of previous results is that low frequency power reflects the construction of linguistic structures (Ding et al., 2016; Kaufeld et al., 2020; Keitel et al., 2018) whereas low frequency phase coherence reflects parsing and segmenting of speech signals (Doelling et al., 2014; Howard & Poeppel, 2010; Kösem & van Wassenhove, 2017; Luo & Poeppel, 2007; Peelle et al., 2013). However, the format in which the brain represents higher-level linguistic structures remains unknown, and novel insights into this issue could have substantial implications for theories of speech comprehension.

### The current study

We investigated whether low frequency neural oscillations reflect differences in syntactic structure. In order to increase the likelihood that any observed patterns are due to representing and processing syntactic structure, we strictly controlled the physical and semantic features of our materials. We extend the work of Ding et al. (2016) and others to ask whether the 1 Hz neural response can be decomposed to reflect separate syntactic structures (phrases versus sentences). To assess this, we used two types of natural speech stimuli in Dutch, namely determiner phrases such as *De rode vaas (The red vase)* and sentences such as *De vaas is rood (The vase is red)*, which combine a subject with a verb into a proposition. Phrases and sentences were matched in the number of syllables (4 syllables), the semantic components (same color and object), the duration in time (1-second, sampling rate 44.1 k Hz), and the overall energy (Root Mean Squared value equals -16 dB).

We formulated a general hypothesis that low frequency neural oscillations would be sensitive to the difference in syntactic structure of the phrases and sentences. However, we did not limit our analysis to low frequency power and phase, as in previous research (Brennan & Martin, 2020; Ding et al., 2016; Kaufeld et al., 2020; Keitel et al., 2018). We hypothesized that the neural response difference between phrases and sentences may manifest itself in a number of dimensions, dimensions that are outside of the view of typical analyses of low frequency power and phase.

We therefore employed additional methods to decompose the neural response to phrases and sentences, to address the following five questions:

#### Question 1

Do phrases and sentences have different effects on brain dynamics as reflected at the functional neural network level (viz., functional connectivity). Neuroscience is rapidly growing interest in investigating functional connectivity to study whole brain dynamics in sensor space (Cabral, Kringelbach, & Deco, 2014; Cohen, 2014, 2015; Hutchison et al., 2013; Sporns, 2010), which can reveal temporal synchronization (viz., phase coherence) between brain regions. Neurophysiological techniques such as EEG and MEG have a high temporal resolution and are suitable for calculating synchronization across frequency bands in functional brain networks (Stam, Nolte, & Daffertshofer, 2007). Describing the temporal synchronization of the neural activity over the whole brain is the first step in decomposing neural responses to high-level variables like syntactic structure. We therefore investigated whether phrases and sentences have different effects on inter-site phase coherence (ISPC), which is considered to reflect the temporal synchronization of neural activity (Cohen, 2014; Lachaux et al., 2000; Mormann, Lehnertz, David, & Elger, 2000).

#### Question 2

Do phrases and sentences differ in the intensity with which they engage connected brain regions? Power connectivity (Cohen, 2011, 2014, 2015) can be used to describe a functional neural network by the energy that is engaged in a cognitive task. Power connectivity is a measure of how different underlying brain regions are connected via the intensity of induced neural responses in the time-frequency space. Differences in power connectivity would imply that phrases and sentences differently impact the distribution and intensity of neural networks involved in speech comprehension.

#### Question 3

Do phrases and sentences have different effects on the coupling between lower and higher frequency activity? This question is related to the theoretical model by Giraud and Poeppel (2012) of a generalized neural mechanism for speech perception below the level of the word. The model, which is focused on syllable-level processing, suggests that presentation of the speech stimulus first entrains an inherent neural response at low frequencies (less than 8 Hz) in order to track to the speech envelope, from which the neural representation of syllables is then constructed. Then, the low frequency neural response evokes a neural response at a higher-frequency (25 to 35 Hz), which reflect the brain’s analysis of phonemic level information. The model proposes coupling between the low and the high frequency neural responses (theta and gamma, respectively) as the fundamental neural mechanism for speech perception up to the syllable. We therefore investigated whether theta-gamma frequency coupling may also differentiate higher-level linguistic structure, namely phrases and sentences.

#### Question 4

Do phrases and sentences also differentially impact neural activity at higher frequencies such as the alpha band? The functional role of alpha band oscillations in perception and memory processing is currently widely debated in systems neuroscience. However, whereas the role for low frequency neural activity in language processing is beyond doubt, whether alpa band activity has an important contribution is not yet agreed upon. Alpha band activity correlates with verbal working memory (Haegens, Osipova, Oostenveld, & Jensen, 2010; Obleser, Wöstmann, Hellbernd, Wilsch, & Maess, 2012; Ten Oever, De Weerd, & Sack, 2020; Wilsch & Obleser, 2016) and auditory attention (Strauß, Wöstmann, & Obleser, 2014; Wöstmann, Herrmann, Maess, & Obleser, 2016; Wöstmann, Herrmann, Wilsch, & Obleser, 2015; Wöstmann, Lim, & Obleser, 2017). Some oscillator models of speech perception consider the induced neural response at alpha as a “top-down” gating control signal (Ghitza, Giraud, & Poeppel, 2013; Giraud & Poeppel, 2012), reflecting domain-general aspects of perceptual processing that are not specific to language processing. Other researchers argue that alpha activity reflects speech intelligibility (Becker, Pefkou, Michel, & Hervais-Adelman, 2013; Dimitrijevic, Smith, Kadis, & Moore, 2017; Obleser & Weisz, 2012), which opens up a potential role for alpha band oscillations in syntactic or semantic processing. We therefore investigated whether phrases and sentences elicited differences in alpha band activity.

#### Question 5

Can we obtain evidence for the differential encoding of phrases and sentences after modelling out physical differences? Neural responses to phrases and sentences comprise a mixture of processes associated with linguistic structure building and with processing acoustic differences. To deal with this issue, one can model which aspects of the neural response encode acoustic differences, and then detect differences between phrases and sentences in the remainder of the neural activity, from which acoustic differences have been regressed out. Previous research using this approach, the spectral temporal response function (STRF), shows that low frequency neural responses robustly represent the acoustic features in the speech (Ding & Simon, 2012a, 2012b, 2013) and that phonemic level processing is reflected in the low frequency entrainment to speech (Di Liberto, O’Sullivan, & Lalor, 2015; Donhauser & Baillet, 2020; Weissbart, Kandylaki, & Reichenbach, 2020). We therefore used the STRF to investigates which dimensions of the neural responses reflect differences between phrases and sentences.

In sum, we investigated different dimensions of the electroencephalography (EEG) response to spoken phrases and sentences. Observing differences between phrases and sentences would serve as a trailmarker on the path towards a theory of the neural computations underlying of syntactic structure formation.

## Results

### Low frequency phase coherence distinguishes phrases and sentences

To answer our first question, whether the low frequency neural oscillations distinguish phrases and sentences, we calculated the phase coherence (for details see Methods). We then performed non-parametric cluster-based permutation tests (1000 permutations) on a 1200-ms time window starting at the audio onset and over the frequencies from 1 Hz to 8 Hz. The comparison indicated that phase coherence was significantly higher for sentences than phrases (p < 1e-4, two-tailed). In the selected latency and frequency range, the effect was most pronounced at central electrodes.

**Fig 2a** shows the temporal evolution, in steps of 50 ms, of this condition effect which is computed as the phase coherence of the phrase condition minus the phrase coherence of the sentence condition. **Fig 2b** shows the time-frequency plot using all the sensors in this cluster, in which the upper and lower panel are the plot for the phrase condition and the sentence condition, respectively.

**Fig 2.**
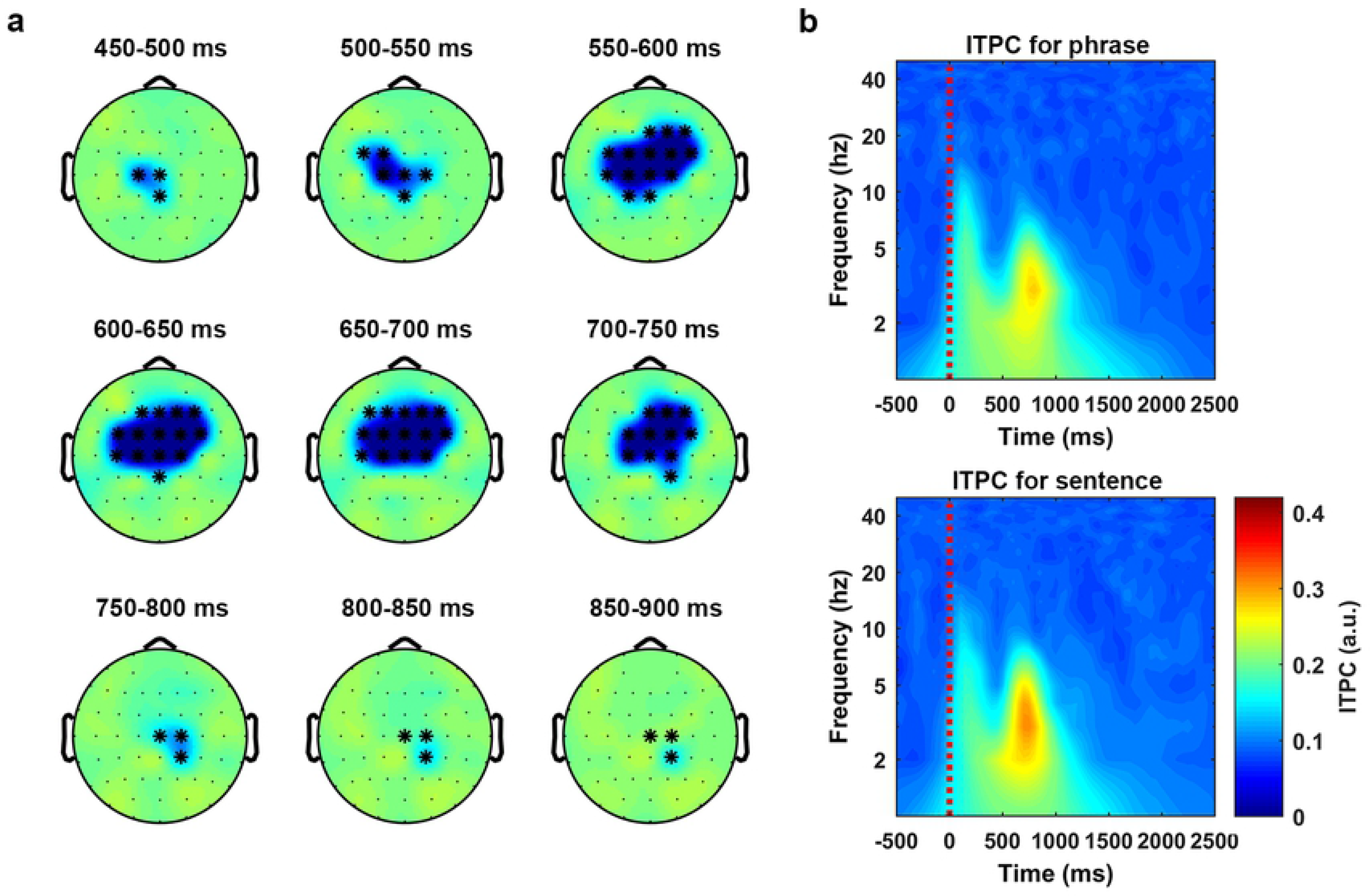
Results of phase coherence. Statistical analysis for comparing the phase coherence (ITPC) difference between phrases and sentences was conducted by using the non-parametric cluster based permutation test (1000 times) on a 1200-ms time window starting at the audio onset and over the frequencies from 2 Hz to 8 Hz. The results indicated that the phase coherence was higher for the sentences than the phrases (p < 1e-4 ***, two-tailed). **(a)** The temporal evolution of the cluster that corresponds to the condition difference between phrases and sentences. The activity was drew by using the ITPC of the phrase condition minus the ITPC of sentence condition. The topographies were plotted in steps of 50 ms. **(b)** ITPC averaged over all the sensors in this cluster. The upper panel and the lower panel shows the ITPC of the phrase condition and the sentence condition, respectively.

The results indicated that the low frequency phase coherence could reliably distinguish phrases and sentences, consistent with the hypothesis that low frequency phase coherence represents cortical computations over speech stimuli (Brennan & Martin, 2020; Doelling et al., 2014; Howard & Poeppel, 2010; Kaufeld et al., 2020; Luo & Poeppel, 2007; Martin, 2016, 2020; Meyer & Gumbert, 2018; Peelle et al., 2013; Rimmele, Morillon, Poeppel, & Arnal, 2018). Our findings therefore suggest that low frequency phase coherence is involved in speech comprehension at the level of syntactic processing.

### Low frequency (< ∼ 2 Hz) phase connectivity degree over the sensor space separates phrases and sentences

We initially calculated phase connectivity over the sensor space by ISPC at each time-frequency bin (for details see Methods section). We then used a statistical threshold method to transform each connectivity representation to a super-threshold count at each bin. After baseline correction, we conducted a 1000 time cluster-based permutation test on a 3500-ms time window starting at the audio onset and over the frequencies from 1 Hz to 8 Hz to compare the degree of the phase connectivity between phrases and sentences (for details see Methods). Phrases and sentences showed a significant difference in connectivity (p < 0.01, two-tailed). The effect corresponded to a cluster extended from ∼1800 ms to ∼2600 ms after the speech stimulus onset, and was mainly located at a very low frequency range (< ∼2 Hz). In the selected latency and frequency range, the effect was most pronounced at the right posterior region.

**Fig 3a** shows the temporal evolution of the condition effect, which is represented by the connectivity degree of the phrase condition minus the connectivity degree of sentence condition (in steps of 100 ms). **Fig 3b** shows the time-frequency plot of the phase connectivity degree, which is averaged across all sensors in this cluster. The left and right panel are the time-frequency plot for the phrase condition and the sentence condition, respectively.

**Fig 3.**
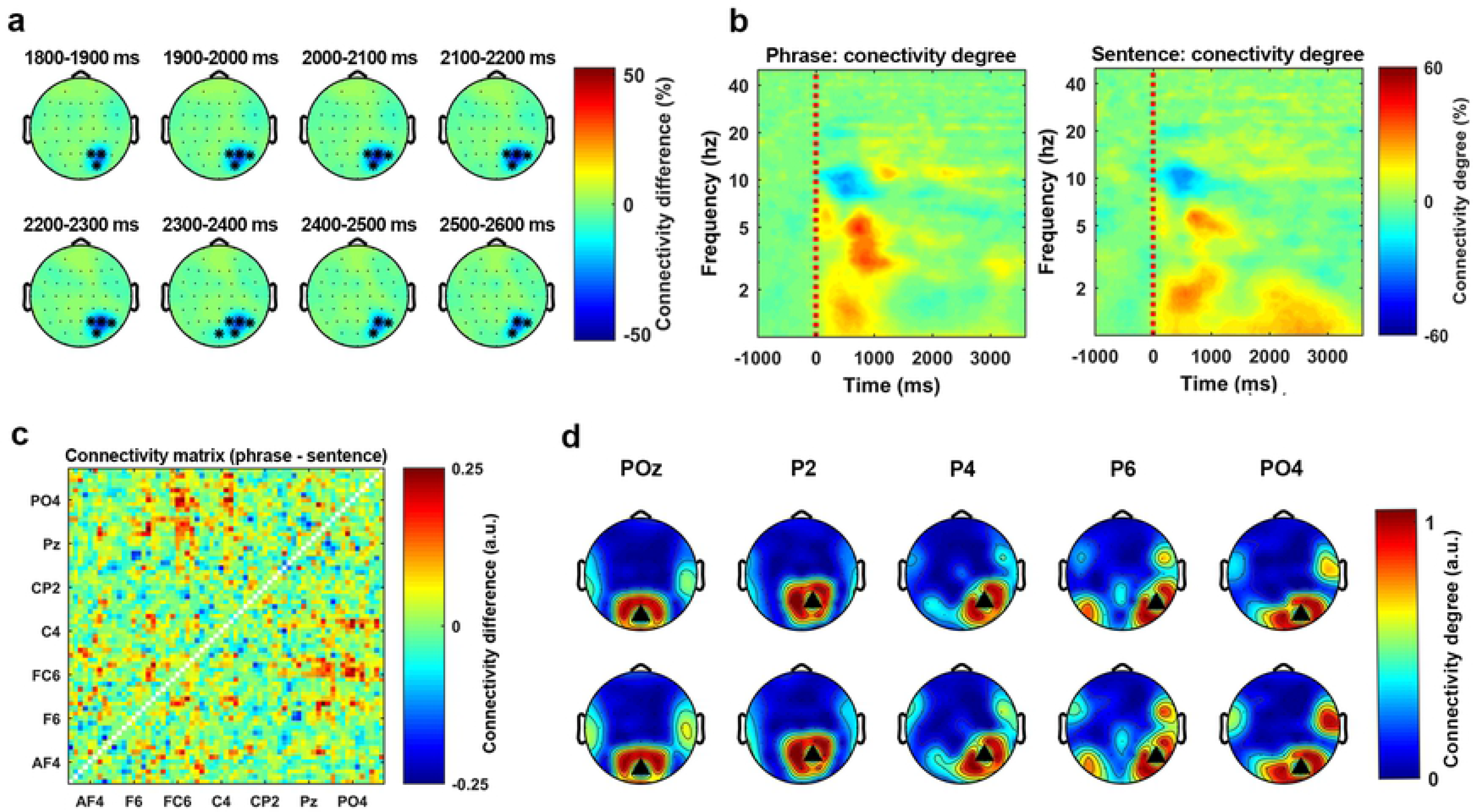
Results of phase connectivity. Statistical analysis for comparing the difference of the phase connectivity degree between phrases and sentences was conducted by using the non-parametric cluster based permutation test (1000 times) on a 3500-ms time window starting at the audio onset and over the frequencies from 1 Hz to 8 Hz. The results indicated that the phase connectivity degree was higher for the sentences than the phrases (p < 0.01**, two-tailed). **(a)** The temporal evolution of the cluster that corresponds to the condition effect. The activity was drew by using the connectivity level of the phrase condition minus the connectivity level of the sentence condition. The topographies were plotted in steps of 100 ms. **(b)** The time-frequency plot of the connectivity degree, which was averaged over all the sensors in this cluster. The left figure and the right figure shows the connectivity level of the phrase condition and the sentence condition, respectively. **(c)** The graph (matrix) level representation of the difference of the phase connectivity. The figure was plotted by using the averaged graph of the phrase condition minus the averaged graph of the sentence condition. **(d)** All the sensors in this cluster were used as the seed sensors to plot the topographical level representation of the phase connectivity. The upper panel and the lower panel shows the phase connectivity of the phrase condition and the sentence condition, respectively.

Since the statistical analysis indicated a phase connectivity degree difference between phrases and sentences, we assessed how this effect was distributed in the sensor space. To do so, we extracted all binarized connectivity matrices that corresponded to the time and frequency range of the cluster and averaged all the matrices in this range for both conditions (details see Methods). **Fig 3c** shows the averaged matrix representation of the sentence condition minus the averaged matrix representation of the phrase condition. This result suggests that the connectivity difference was mainly localized at the frontal-central area. After extracting the matrix representation, we used all sensors of this cluster as seeds to plot connectivity topographies for both conditions. **Fig 3d** shows the pattern of the thresholded phase connectivity. The black triangles represent the seed sensors. The upper panel and lower panel represent the phrase and the sentence condition, respectively. The figure shows how the phase connectivity (synchronization) is distributed on the scalp in each condition. From this figure we can see that the overall degree of the phase connectivity was stronger for the sentence condition than the phrase condition.

The analysis indicated that the phase connectivity degree over the sensor space at the low frequency range (< ∼ 2 Hz) could reliably separate the two syntactically different stimuli and that the effect was most prominent at the right posterior region.

### Phase Amplitude Coupling (PAC) as a generalized neural mechanism for speech perception

To assess whether PAC distinguished phrases and sentences, we calculated the PAC value at each phase-amplitude bin for each condition, and then transformed it to the PAC-Z (for details see Methods). The grand average (average over sensors, conditions and participants) of the PAC-Z showed a strong activation over a region from 4 Hz to 10 Hz for the frequency of phase response and from 15 Hz to 40 Hz for frequency of the amplitude. We, therefore, used the averaged PAC-Z value in this region as the index for sensors clustering. For each participant, we first selected 8 sensors that had the highest PAC-Z (conditions averaged) at each hemisphere. Averaging over sensors was conducted separately for the conditions (phrase and sentence) and the two hemispheres (see **Fig 4a**). The Bonferroni correction was performed to address the multiple comparison problem. This resulted in the z-score of 3.73 for p<0.05 (the z-score corresponded to the p-value equals 0.05 divided 11 (the number of phase bins)*12(the number of amplitude bins)*4(the number of conditions)). From the results, we can see that there was a strong low frequency phase (4 Hz to 10 Hz) response entrained to high frequency amplitude (15 Hz to 40 Hz). The results indicate that the PAC was introduced when participants listened to the speech stimuli.

**Fig 4.**
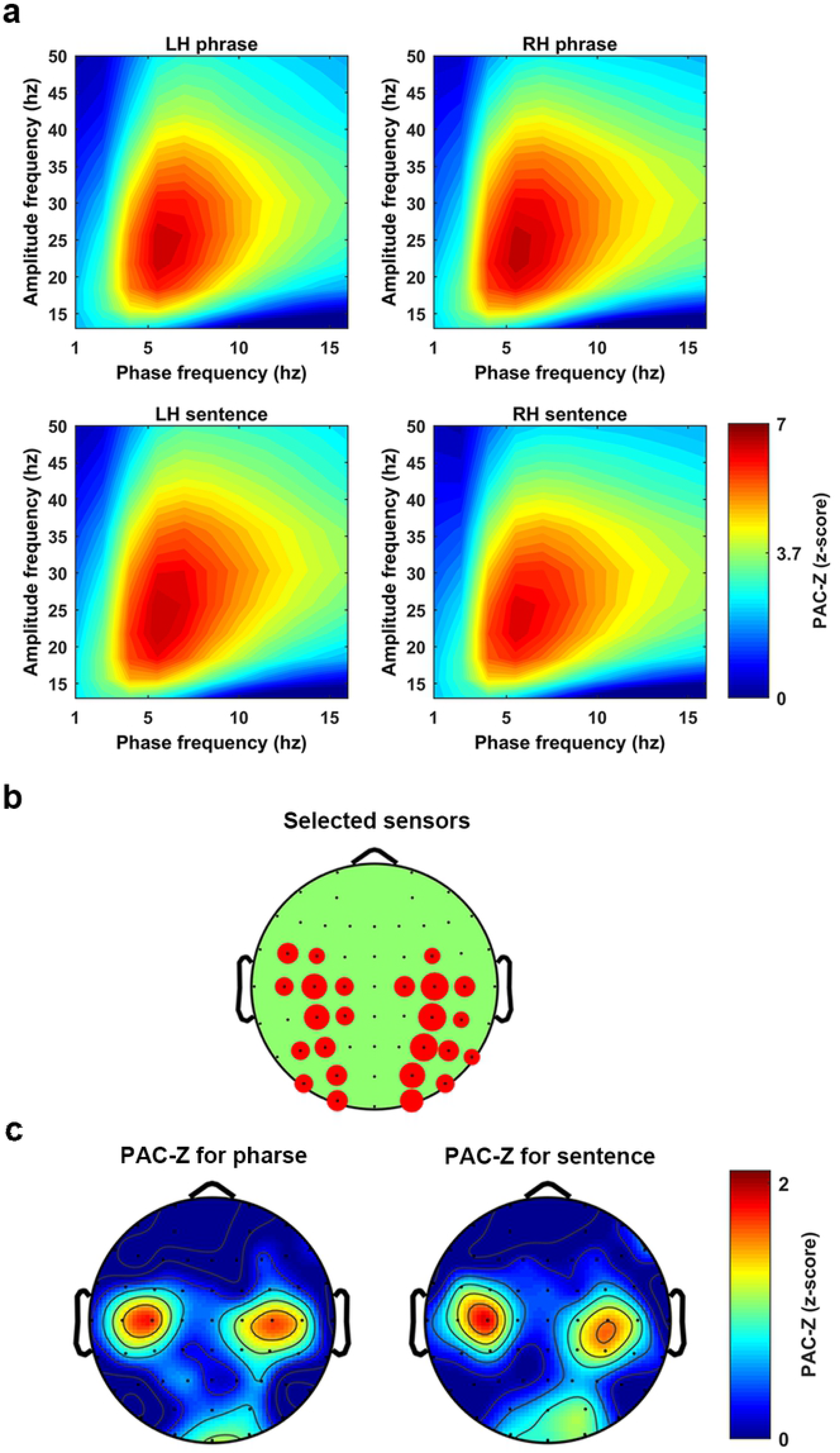
Results of phase amplitude coupling. The z-score transformed phase amplitude coupling, PAC-Z. **(a)** The PAC-Z for the phrase condition and the sentence condition at each hemisphere. Each figure was created by averaging 8 sensors which showed the biggest PAC-Z over the region which defined by the phase frequency from 4 to 10 Hz and the amplitude frequency from 15 to 40 Hz. A z-score transformation with Bonfferroni correction was conducted to test the significance, which lead to the threshold to be 3.73 corresponding to p=0.05. **(b)** The figure shows how sensors were selected at each hemisphere. The bigger the red circle indicates the more times this sensor was selected. **(c)** The topographical distribution of the PAC-Z. The figure indicates that the PAC was largely localized at the bilateral central areas.

**Fig 4b** shows how the sensors were selected. The larger the red circle indicates the more often the sensor was selected across participants.

**Fig 4c** shows the topographical representation of the PAC-Z. The activity in these figures was the averaged PAC-Z values over the frequency range for phase (4 to 10 Hz) and the frequency range for amplitude (15 to 40 Hz). The results indicate that when the participants listened to the speech stimuli, PAC was introduced symmetrically at both hemispheres over the central area. This could be evidence for the existence of PAC when speech stimuli are being processed. However, both the parametric and the non-parametric statistical analysis failed to show a significant difference of the PAC-Z between phrases and sentences, which means we do not have evidence to show that the PAC was related to syntactic information processing. Therefore, our results suggest the PAC could be a generalized neural mechanism for speech perception, rather than a mechanism specifically recruited during the processing of higher-level linguistic structures.

### Alpha band inhibition reflects syntactic structure

To query whether neural oscillations at alpha band reflect the processesing of syntactic structure, we calculated the induced power. The grand average (over all participants and all conditions) of the induced power showed a strong inhibition at the alpha band (∼7.5 to 13.5 Hz). Therefore, we checked whether this alpha band inhibition could separate the two types of linguistic structures. Statistical analysis was conducted using the nonparametric cluster-based permutation test (1000 times) over the frequencies of alpha band in the 1000 ms time window from the audio onset for details see Methods). The results indicated that the alpha band inhibition was stronger for the phrase condition than the sentence condition (p < 0.01, two-tailed). In the selected time and frequency range, this effect corresponded to a cluster that lasted from ∼350 ms to ∼1000 ms after the audio onset and was largely localized at the left hemisphere, though the right frontal-central sensors were also involved during the temporal evolution of this cluster. **Fig 5a** shows the temporal evolution of this cluster in steps of 50 ms using the induced power of the phrase condition minus the induced power of the sentence condition. **Fig 5b** shows the time-frequency plot of the induced power using the average of all the sensors in this cluster. The upper and lower panel shows the phrase condition and the sentences condition, respectively. From these figures, we can see that the alpha band inhibition was stronger for the phrase condition than the sentence condition. These results show that the processing of phrases and sentences is reflected in the intensity of the induced neural response in the alpha band.

**Fig 5.**
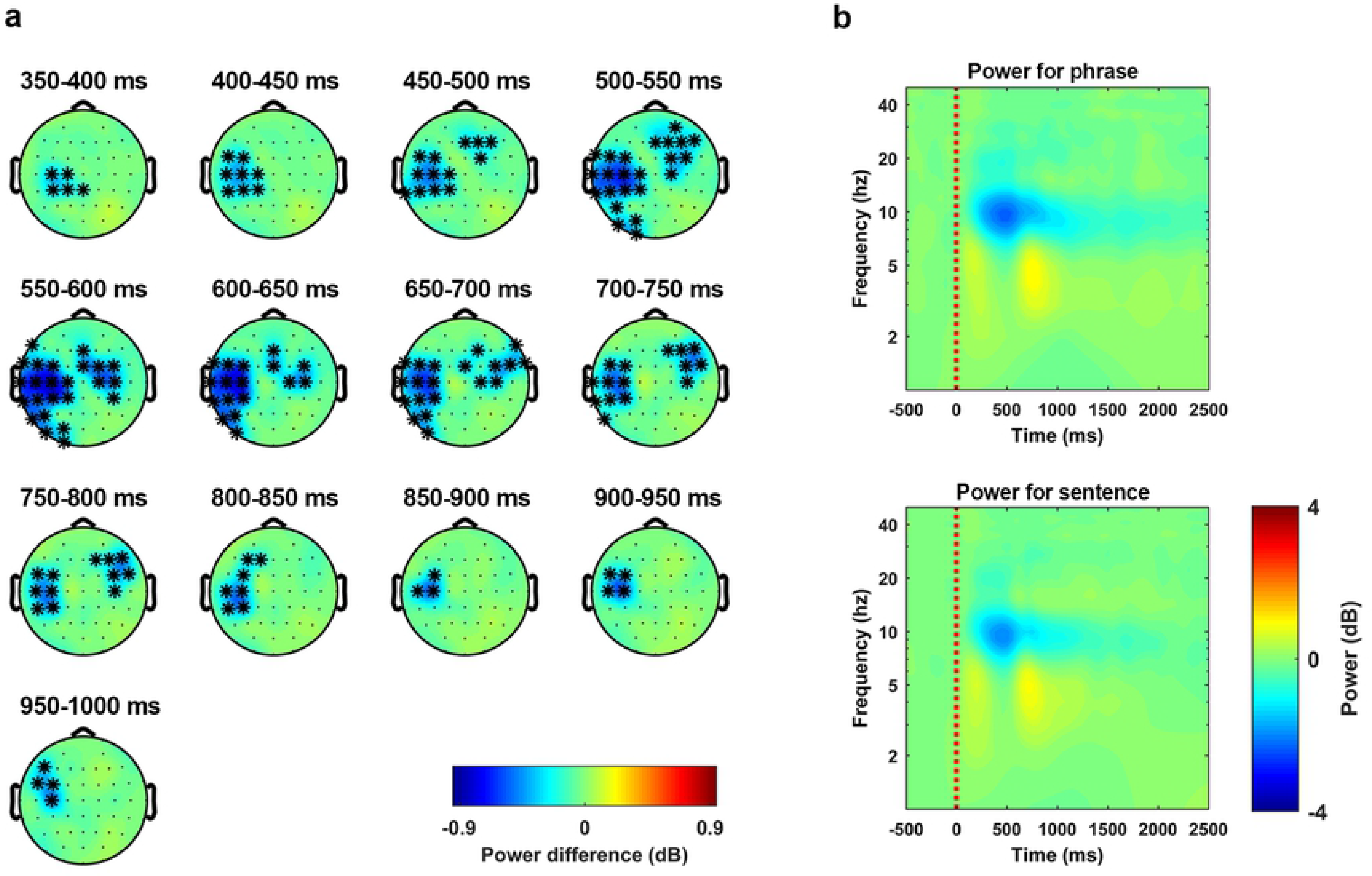
Results of induced neural response. Statistical analysis for comparing the induced power difference between phrases and sentences was conducted by using the non-parametric cluster based permutation test (1000 times) on a 1000-ms time window starting at the audio onset and over the frequencies from 7.5 Hz to 13.5 Hz. The results indicated that the phase coherence was higher for sentences than phrases (p < 0.01 **, two-tailed). **(a)** The temporal evolution of the cluster that corresponds to the condition difference between phrases and sentences. The activity was drew by using the induced power of the phrase condition minus the induced power of sentence condition. The topographies were plotted in steps of 50 ms. **(b)** Induced power averaged over all the sensors in this cluster. The upper panel and the lower panel shows the induced power of the phrase condition and the sentence condition, respectively.

### The power connectivity degree in the alpha band indicates network level separation of phrases and sentences

We calculated power connectivity in each sensor-pair at each time-frequency bin using Rank Correlation (for details see Methods). The grand average (over all participants and all conditions) of the power connectivity level showed a strong inhibition at the alpha band from 100 ms to 2200 ms after the audio onset. This region, which showed a strong power connectivity inhibition, was defined as the Region of Interest (ROI). For each participant, we selected eight sensors at each hemisphere that showed the biggest inhibition of the condition averaged power connectivity at the ROI. Averaging across all the selected sensors was followed up, which resulted in four conditions for each participant (left-phrase, left-sentence, right-phrase and right-sentence).

**Fig 6a** shows the power connectivity degree which was averaged over all participants for each condition. To check whether power connectivity degree separated the phrases and the sentences, a Stimulus-Type*Hemisphere 2-way repeated measure ANOVA was conducted. The comparison revealed a main effect of Stimulus-Type (F (1, 14) = 5.28, p = 0.033). Planned post hoc comparison using paired sample t-tests on the main effect of the Stimulus-Type showed that the power connectivity inhibition was stronger for the phrases than the sentences (t(29) = 2.82, p = 0.0085). **Fig 6b** shows the results of the comparison. **Fig 6c** shows how sensors were selected. The bigger the red circle indicates the more times the sensor was selected.

**Fig 6.**
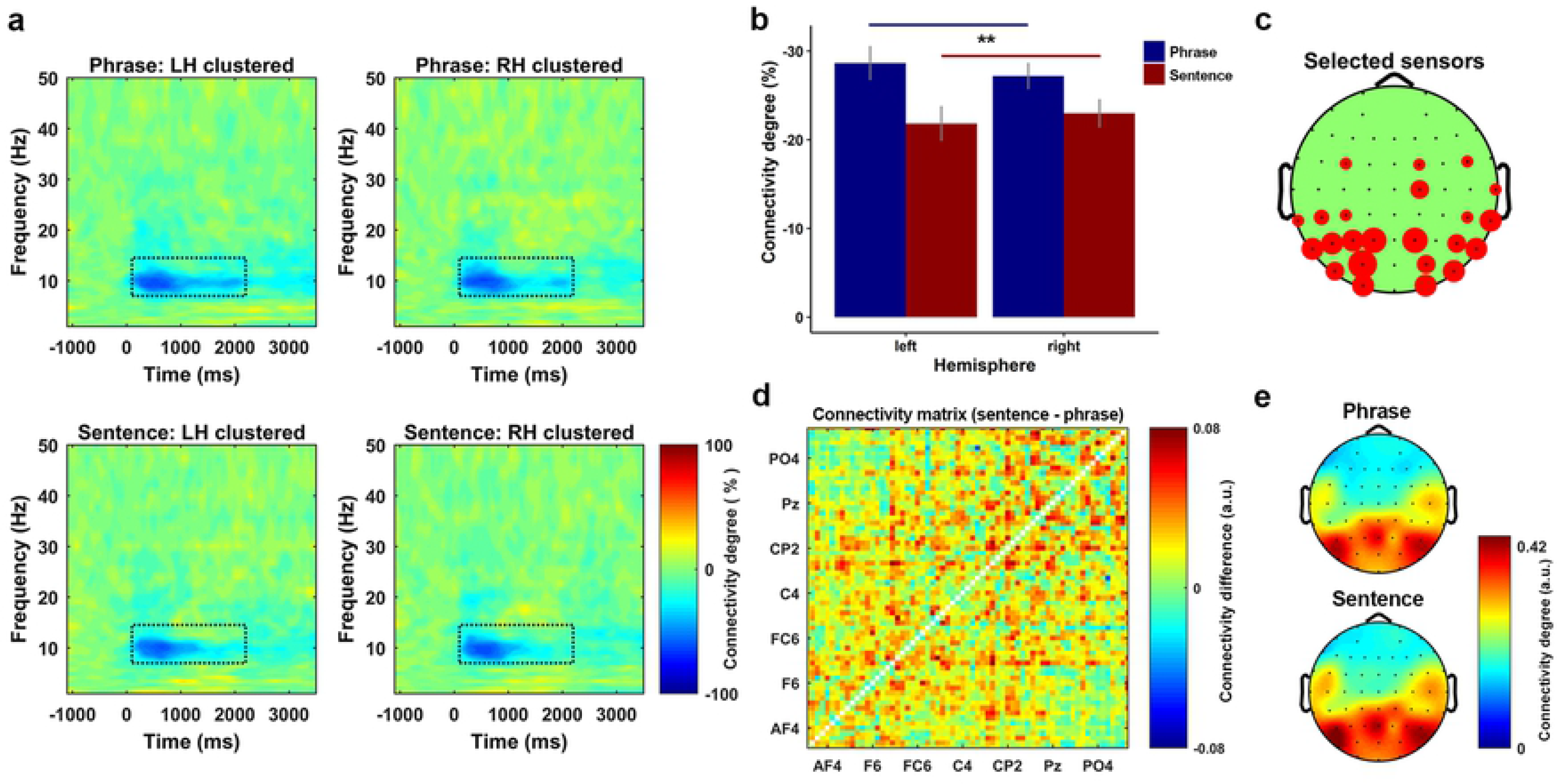
Results of power connectivity. **(a)** Power connectivity degree for all conditions. The left and right figure of the upper and lower panel shows the power connectivity degree for the phrase condition and the sentence condition at the left and right hemisphere, respectively. Each plot was clustered by the sensors at each hemisphere that showed the biggest inhibition level of the grand averaged power connectivity. **(b)** The results of two-way repeated measure ANOVA of power connectivity on the factors of stimulus-type (phrase or sentence) and hemisphere (left or right) indicated a significant main effect of stimulus-type, post hoc comparison on the main effect indicated that the overall inhibition level of the power connectivity was stronger for the phrases than the sentences (t(29) = 2.82, p = 0.0085 **, two-sided). **(c)** How sensors were selected for the clustering. The bigger the red circle indicates the more times the sensor was selected across participants. **(d)** The connectivity differences between the phrases and the sentences on all sensor-pair. The plot using the average of the binarized connectivity matrix of the sentence condition minus the phrase condition. The figure shows that the connectivity degree over the sensor space for the sentence condition was higher than the phrase condition. **(e)** Topographical plot of the binarized connectivity that was clustered by the sensors showing biggest inhibition level of the power connectivity. The upper and lower panel shows the phrase and sentence condition, respectively.

Since the degree of the power connectivity over the alpha band indicated a separation between the phrases and the sentences, we also checked how this difference was distributed in the sensor space. To do so, we extracted the binarized power connectivity representations (matrices) that are located in the ROI, then averaging was performed for each condition across all connectivity matrices. **Fig 6d** shows the difference of the power connectivity degree over the sensor space using the average of the binarized sentence connectivity matrix minus the average of the binarized phrase connectivity matrix. The results indicate that the inhibition of the power connectivity was stronger for phrases than for sentences. In other words, the overall level of the power connectivity was higher for sentences than for phrases. **Fig 6e** is the topographical representation of the power connectivity, which was plotted using the binarized power connectivity of the selected sensors. The upper and lower panel are the phrase condition and the sentence condition, respectively. From this figure, we can see that the difference was largely localized at the bilateral central area, and more strongly present at the left than the right hemisphere. These results reflect that the neural network which was organized by the intensity of the induced power at alpha band was different for the two syntactic structures.

### Different encoding states for phrases versus sentences in both temporal and spectral dimensions

Previous research has shown that the low frequency neural response reliably reflect the phase locked encoding of the acoustic features of speech (Ding & Simon, 2012a, 2012b). Therefore, we initially tested whether the neural response from all canonical frequency bands could equally reflect the encoding of the acoustic features. To do so, we fitted the STRF for each condition at all frequency bands, which are Delta (< 4 Hz), Theta (4 to 7 Hz), Alpha (8 to 13 Hz), Beta (14 to 30 Hz) and Low-Gamma (31 to 50 Hz), respectively. Then we compared the real performance of the STRFs to the random performance of them (for details see Methods). **Fig 7a** shows the results of this comparison. The blue and red dots represent the real performance of the STRFs, the error bar presents 1 s.e.m on each side. The small gray dots represents the random performance (1000 times in each frequency band per condition). The upper boarder, which is delineated by these gray dots represents the 97.5 percentiles of the random performance. The performance of the STRFs was above chance level only at the low frequency (Delta and Theta) band, which was consistent with previous research (Ding & Simon, 2012a, 2012b). Our results verified that the low frequency STRF reliably reflected the relationship between the acoustic features of speech and the neural response at low frequencies.

**Fig 7.**
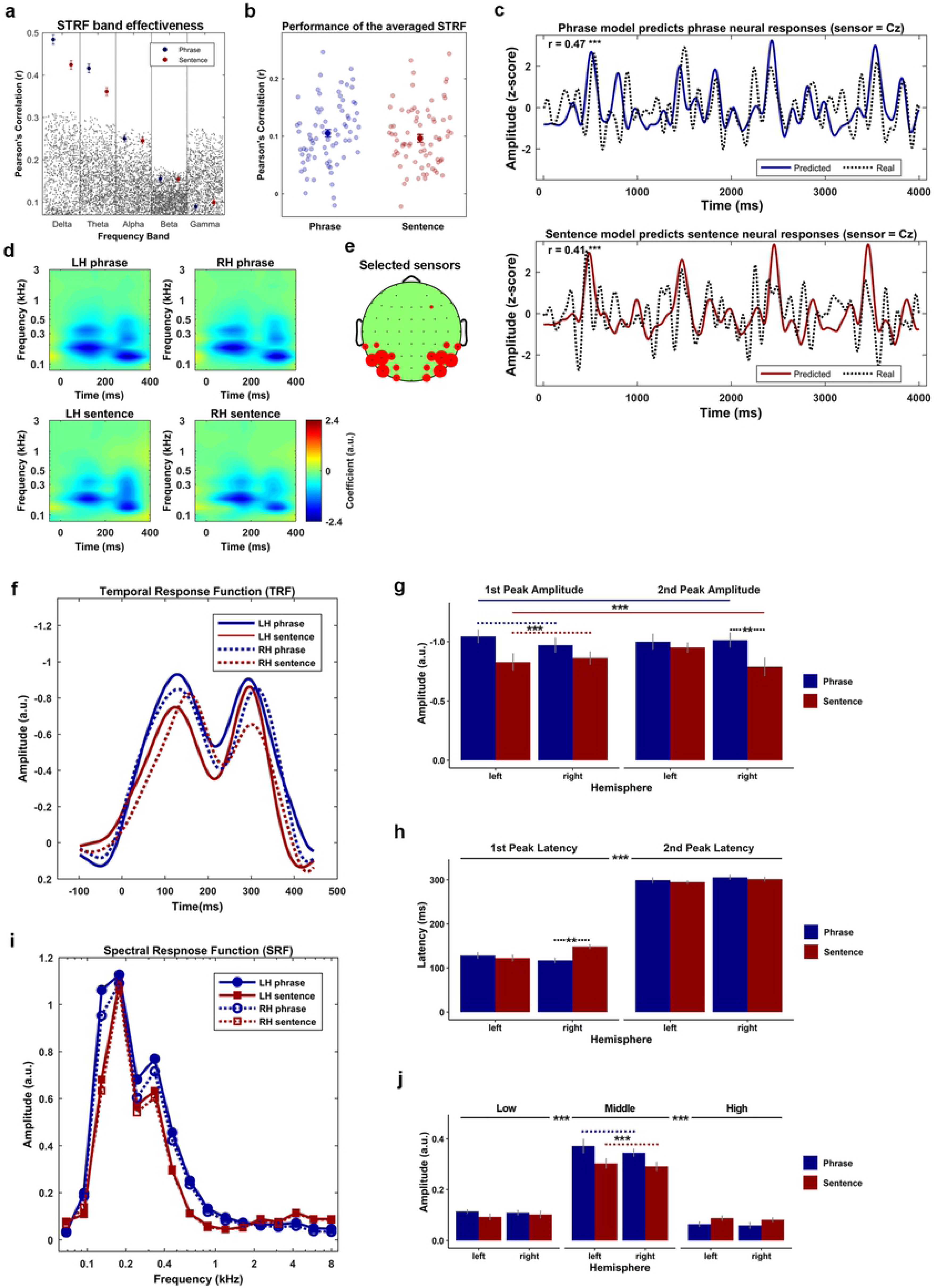
Results of STRF. **(a)** The comparison between the real performance and the random performance of the STRF in each frequency band. The results suggested that only the performance of the STRF in the Delta band (< 4 Hz) and Theta band (4-8 Hz) was better than the random performance. The blue and red dots represent the real performance of the STRFs for the phrase condition and the sentence condition, respectively. The error bar represents two S.E.M across the mean. The gray dots represents the random performance. **(b)** The performance of the low frequency (< 8 Hz) STRF which was averaged across all participants. The solid blue and red dot represents the averaged performance across all testing trials. The error bar represents two S.E.M across the mean. The transparent blue and red dots represent the model’s performance on each testing trial for the phrase condition and the sentence condition, respectively. The results indicate no performance differences of the model between the phrase condition and the sentence condition. **(c)** The comparison between the real neural response (dashed lines) and the averaged model predicted response (solid blue for the phrase, solid red for the sentence) at a sample sensor Cz. The results show that both the averaged model for the phrase condition (r = 0.47 ***, n=1024) and the sentence condition (r=0.41 ***, n=1024) had a good performance. **(d)** STRF clustered by the selected sensors which showed the biggest amplitude (negative) on the time from 0 to 400 ms and on frequency between 0.1 kHz to 0.8 kHz. The figure on the left and right side of the upper panel represent the clustered STRF for the phrases at the left and right hemisphere, respectively. The corresponding position of the lower panel represent the clustered model for the sentence condition. **(e)** The figure shows how the sensors were selected, in which the bigger the red circle represents the more times the sensor was selected across all participants. **(f)** The TRFs that were decomposed from the STRFs, in which the blue and red lines represent the phrase condition and the sentence condition, respectively. The solid and the dashed lines represent the left and right hemisphere, respectively. **(g)** The comparison of the magnitude of the TRFs, the blue and red bars represent the phrase condition and the sentence condition, respectively. The error bar shows 1 s.e.m across the mean on each side. A 3-way repeated measure ANOVA of the peak magnitude was conducted on the factors of stimulus-type (phrase or sentence), Hemisphere (left or right) and peak-type (∼100 ms or ∼300 ms). The results indicated a main effect of stimulus-type and a 3-way interaction. The post hoc comparison on the main effect of stimulus-type suggested that the amplitude (negative) was stronger for the phrase condition than the sentence condition (t (59) = 4.55, P < 2e-5 ***). To investigate the 3-way, Stimulus-type*Peak-type*Hemisphere, interaction, two 2-way repeated measure ANOVA with the Bonferroni correction were conducted on the factors of Hemisphere and Audio-type at each level of the Peak-type. The results indicated a main effect of stimulus-type at the first peak (F (1, 14) = 8.19, p = 0.012 *) and a 2-way Hemisphere*stimulus-type interaction at the second peak (F (1, 14) = 6.42, p = 0.023 *). At the first peak, a post hoc comparison on the main effect of stimulus-type was conducted using a paired sample t tests, the results showed that the magnitude of the phrase condition was higher than the magnitude of the sentence condition (t(29) = 3.49, p = 0.001 ***). For the 2-way, Hemisphere*stimulus-type, interaction at the second peak, the paired sample t tests with Bonferroni correction was conducted to compare the difference of the magnitude between the phrase condition and the sentence condition at each hemisphere. The results indicate that the magnitude at the second peak was stronger for the phrase condition than the sentence condition in the right hemisphere (t (14) = 3.21, p = 0.006 **), but not the left hemisphere (t (14) = 0.86, p = 0.40). **(h)** The comparison of the peak latency of TRFs, the blue and red bars represent the phrase condition and the sentence condition, respectively. The error bar shows 1 s.e.m across the mean on each side. A 3-way repeated measure ANOVA of the peak latency was conducted on the factors of stimulus-type (phrase or sentence), Hemisphere (left or right) and peak-type (∼100 ms or ∼300 ms). The results indicated a main effect of peak-type and a 3-way interaction. The post hoc comparison on the main effect of peak-type suggested that the latency of the first peak was significantly faster than the second peak (t (59) = 38.89, p < 2e-16 ***). The post hoc comparison on the 3-way interaction with the Bonderroni correction on the factors of Hemisphere and stimulus-type for each level of the Peak-type suggested a 2-way Hemisphere*stimulus-type interaction at the first peak (F (1, 14) = 12.83, p = 0.002***). The post hoc comparison on this 2-way interaction using paired sample t tests with the Bonferroni correction indicated that the latency at the first peak was significantly longer for the sentences than the phrases at the right hemisphere (t(14) = 3.55, p = 0.003 ***), but not the left hemisphere (t(14) = 0.58, p = 0.56). **(i)** The SRFs which were decomposed from the STRFs, in which the red and blue lines represent the phrase condition and the sentence condition, respectively, the solid and the dashed lines represent the left and right hemisphere, respectively. **(j)** The comparison of the amplitude of the SRFs. The SRF was first separated into three bands, low ( < 0.1 kHz), middle ( 0.1 to 0.8 kHz) and high ( > 0.8 kHz) based on the averaged frequency response of the STRF, then a 3-way repeated measure ANOVA of the amplitude was conducted on the factors of stimulus-type (phrase or sentence), Hemisphere (left or right) and frequency-band (low, middle and high). The results indicated a main effect of Band-type (F (2, 28) = 119.67, p < 2e-14 ***) and a 2-way, Band-type*stimulus-type, interaction (F (2, 28) = 27.61, p < 3e-7 ***). The post hoc comparison on the main effect of Band-type using paired sample t tests with Bonferroni correction showed that the magnitude of the middle frequency band was stronger than the low frequency band (t(59) = 17.9, p < 4e-25 ***) and high frequency band (t(59) = 18.7, p < 5e-26 ***). The post hoc comparison using paired sample t tests with the Bonferroni correction on the, Band-type*stimulus-type, interaction showed that the amplitude of the SRF was stronger for the phrases than the sentences only at middle frequency band (t (29) = 4.67, p < 6e-5 ***).

Since only low frequency neural responses robustly reflected the encoding of the speech stimuli, we fitted the STRF for both conditions using the neural response that was low passed at 9 Hz. Leave-one-out cross validation was used to maximize the performance of the STRFs (for details see Methods). **Fig 7b** shows the performance of the STRF for each condition. The transparent dots, blue for phrases and red sentences, represent the model’s performance on each testing trial. The solid dots represent the model’s performance that was averaged over all trials, the error bars represent 1 s.e.m on each side of the mean. A paired sample t-test was used to compare the performance between the phrase condition and the sentence condition. No evidence was shown to indicate a performance’s difference between these two conditions (t (74) = 1.25, p = 0.21). The results indicate that the STRF were fitted equally well for phrases and sentences. Thus any difference in temporal-spectral features between the STRF of phrases and sentences cannot be driven by the model’s performance. **Fig 7c** shows the comparison between the real neural response and the model predicted response at the sample sensor Cz. The upper and lower panel shows the performance of the STRF to phrases (r = 0.47, N=1024, p < 1e-5) and sentences (r = 0.41, N=1024, p < 1e-5), respectively.

The grand average of the STRFs was negative from 0 to 400 ms in the time domain and from 100 to 1000 Hz in the frequency domain, and the sensor clustering of the STRF was conducted based on the averaged activation these domain periods. More concretely, we selected eight sensors at each hemisphere for each participant, which showed the strongest averaged magnitude (negative) in this region. **Fig 7d** shows the clustered STRFs that were averaged across all participants. **Fig 7e** shows how the sensors were selected across the participants, in which the bigger the red circle indicates the more times a given sensor was selected.

To compare the differences of the kernel (STRF) in both the temporal and spectral dimensions, the TRF and the SRF were extracted separately for each condition (for details see Methods).

**Fig 7f** shows the TRFs that were averaged across all participants. The grand average of all TRFs showed two peaks at ∼100 ms and ∼300 ms. We therefore defined the first temporal window from 50 to 150 ms (center at 100 ms) and the second temporal window from 250 to 350 (center at 300 ms) for the search for the magnitude and the latency of these two peaks. The latency of each peak was defined as the time when it appeared. The magnitude of each peak was defined as the average magnitude over a 5 ms window on both sides around it. After extracting the magnitude and the latency of these two peaks, a Stimulus-type*Peak-type*Hemisphere 3-way repeated measure ANOVA was conducted on both the magnitude and the latency. For the magnitude of the TRF (**Fig 7g**), the statistical comparison showed a significant main effect of Stimulus-type (F (1, 14) = 13.58, P = 0.002) and a significant 3-way, Stimulus-type*Peak-type*Hemisphere, interaction (F (1, 14) = 15.25, P = 0.001).

The post hoc comparison on the main effect of Stimulus-Type using paired-sample t-tests showed that the magnitude for phrases was significantly stronger than the magnitude for sentences (t(59) = 4.55, P < 2e-5). The results suggest that the instantaneous neural activity in response to phrases had a stronger phase-locked dependency on the acoustic features than in response to sentences.

To investigate the 3-way, Stimulus-type*Peak-type*Hemisphere, interaction, two 2-way repeated measure ANOVA with the Bonferroni correction were conducted on the factors of Hemisphere and Stimulus-type at each level of the Peak-type. The results indicated a main effect of Stimulus-Type at the first peak (F (1, 14) = 8.19, p = 0.012) and a 2-way Hemisphere*Stimulus-Type interaction at the second peak (F (1, 14) = 6.42, p = 0.023).

At the first peak, we conducted a post hoc comparison on the main effect of stimulus-type using a paired sample t tests, which showed that the magnitude of the phrase condition was higher than the magnitude of the sentence condition (t(29) = 3.49, p = 0.001). The results indicate that the neural activity was more strongly driven by the acoustic features that were presented ∼100 ms ago when phrases than when sentences were presented.

For the 2-way, Hemisphere*Stimulus-Type, interaction at the second peak, the paired-sample t-tests with Bonferroni correction was conducted to compare the difference of the magnitude between phrases and sentences at each hemisphere. The results indicate that the magnitude at the second peak was stronger for phrases than sentences in the right hemisphere (t (14) = 3.21, p = 0.006), but not the left hemisphere (t (14) = 0.86, p = 0.40). The findings suggest that, at the right hemisphere, the instantaneous neural activity of the phrases was more strongly driven by the acoustic features that were present ∼300 ms than it was under sentences.

For the latency of the TRF (**Fig 7h**), the comparison showed a main effect of the Peak-type (F (1, 14) = 1e+3, p<1e-14) and a 3-way, Stimulus-type*Peak-type*Hemisphere, interaction (F (1, 14) = 8.04, p = 0.013).

The post hoc comparison for the main effect of the Peak-type with paired sample t tests showed, as expected, that the latency of the first peak was significantly shorter than the second one (t(59) = 38.89, p < 2e-16). The result is clear since regardless of search method for the analysis time windows, the latency of the first one will always shorter than the second one.

To investigate the 3-way, Stimulus-type*Peak-type*Hemisphere, interaction, two 2-way repeated measures ANOVA with the Bonferroni correction were conducted on the factors of Hemisphere and stimulus-type for each level of the Peak-type. The comparison suggested a 2-way Hemisphere*Stimulus-Type interaction at the first peak (F (1, 14) = 12.83, p = 0.002). The post hoc comparison on this 2-way interaction using paired sample t-tests with the Bonferroni correction indicated that the latency at the first peak was significantly longer for sentences than for phrases at the right hemisphere (t(14) = 3.55, p = 0.003), but not the left hemisphere (t(14) = 0.58, p = 0.56). The results suggest that, within the first temporal window (∼50 to 150 ms), only at the right hemisphere, the neural response of the sentences was dominantly driven by the acoustic features earlier in time than the acoustic features that drove the neural response of the phrases.

**Fig 7i** shows the SRFs that were averaged across all participants. The grand average of the STRFs suggested that the activation of the kernel was most prominent in the frequency range from 0.1 kHz to 0.8 kHz. To compare the differences of the encoding of the acoustic features in the spectral dimension, we separated the SRF into three frequency bands, which were lower than 0.1 kHz, 0.1 to 0.8 kHz and higher than 0.8 kHz. We then averaged the response in each extracted frequency band for each condition. The statistical comparison was conducted using 3-way repeated measure ANOVA on the factors of Hemisphere, stimulus-type and Band-type. The results (**Fig 7j**) indicated a main effect of Band-type (F (2, 28) = 119.67, p < 2e-14) and a 2-way, Band-type*stimulus-type, interaction (F (2, 28) = 27.61, p < 3e-7).

The post hoc comparison on the main effect of Band-type using paired-sample t-tests with Bonferroni correction showed that the magnitude of the middle frequency band was stronger than that of the low frequency band (t(59) = 17.9, p < 4e-25) and high frequency band (t(59) = 18.7, p < 5e-26). The results indicate that the acoustic features from different frequency bands contributed differently to the evoked neural response. In other words, for both conditions, the neural response was dominantly driven by the encoding of the acoustic features from 0.1 kHz to 0.8 kHz, which was considered as the spectral-temporal features at the range of the first formant (Catford, 1988; Jeans, 1968; Titze et al., 2015; Titze & Martin, 1998).

The post hoc comparison using paired-sample t-tests with the Bonferroni correction on the, Band-type*Stimulus-Type, interaction showed that the amplitude of the SRF was stronger for the phrase condition than the sentence condition only at the middle frequency band (t (29) = 4.67, p < 6e-5). The results indicate that at the middle frequency range, the neural response of phrases was more strongly predicted solely by modelling the encoding of the acoustic features than it was in the sentence condition. This pattern of results indicates that the neural representation of sentences is more abstracted away from the neural response that is driven by the physicality of the stimulus.

## Methods

### Participants

Fifteen right-handed Dutch native speakers, 22-35 years old, 7 males, participated in the study. All participants were undergraduate or graduate students. Participants reported no history of hearing impairment or neurological disorder and were paid for their participation. The experimental procedure was approved by the Ethics Committee of the Social Sciences Department of Radboud University. Written informed consent was obtained from each participant before the experiment.

### Stimuli

We selected fifty line-drawings of common objects from a standardized corpus (Snodgrass & Vanderwart, 1980). The Dutch names of all objects were monosyllabic and had non-neuter lexical gender. In our experiment, the objects appeared as colored line-drawing on a grey background. More specifically, we presented each line-drawing in five colors: blue (blauw), red (rood) yellow (geel), green (groen) and purple (paars). In total, this yielded 250 pictures. The line-drawings were sized to fit into a virtual frame of 4 cm by 4 cm, corresponding to 2.29° of visual angle for the participant.

We then selected ten figures with different objects in each color, without object replacement between colors, to create speech stimuli. For each selected line-drawing, a 4-syllable, phrase-sentence pair was created, i.e. *De rode vaas (The red vase) and De vaas is rood (The vase is red).* In total, we had 100 speech stimuli (50 phrases and 50 sentences). All stimuli were synthesized by a Dutch male voice, “Guus”, of an online synthesizer (www.readspeaker.com). All speech stimuli were 733 ms to 1125 ms in duration (Mean= 839 ms, SD=65 ms). In order to normalize the synthesized auditory stimuli, they were first resampled to 44.1 kHz. Then all speech stimuli were adjusted by truncation or zero padding at both ends to 1000 ms without missing any meaningful dynamics. Then 10% at both ends of each stimulus was smoothed by a linear ramp (a sine wave) for removing the abrupt sound burst. Finally, for normalizing the intensity of speech stimuli, the root mean square (RMS) value of each stimulus was normalized to -16 dB.

### Experimental Procedure

Each trial started by a fixation cross at the screen center (500 ms in duration). Participants were asked to look at the fixation. Immediately after the fixation cross had disappeared, the participants heard a 1000 ms long spoken stimulus, either a phrase or a sentence, followed by three-second silence; then the participants were asked to perform one of three types of task, indicated to them by an index (1, 2 or 3 showing at the screen center, 500 ms in duration). If the index was ‘1’, they did a linguistic structure discrimination task (***type-one task***), in which they had to judge whether the spoken stimulus was a phrase or a sentence and indicate their judgment by pressing a button. If the index was ‘2’, a picture would follow (200 ms in duration) and would be shown after a 1000 ms gap after the index number. Then participants would do a color matching task (***type-two task***), in which they had to judge whether the color described in the spoken stimulus matches the color of the shown picture and indicate their judgment by pressing a button. If the index is ‘3’, they would experience the same procedure as the type two task, except for they would do an object matching task (***type-three task***), judging the matching of object between the spoken stimulus and the picture and indicating their respond by pressing a button. All responses were recorded via a parallel port response box, in which each one of the two buttons were represented as phrase/match and sentence/mismatch, respectively. Each response was followed by a silent interval of 3 to 5.2 seconds (jittered). The experimental procedure is depicted in the **Fig 8**.

**Fig 8.**
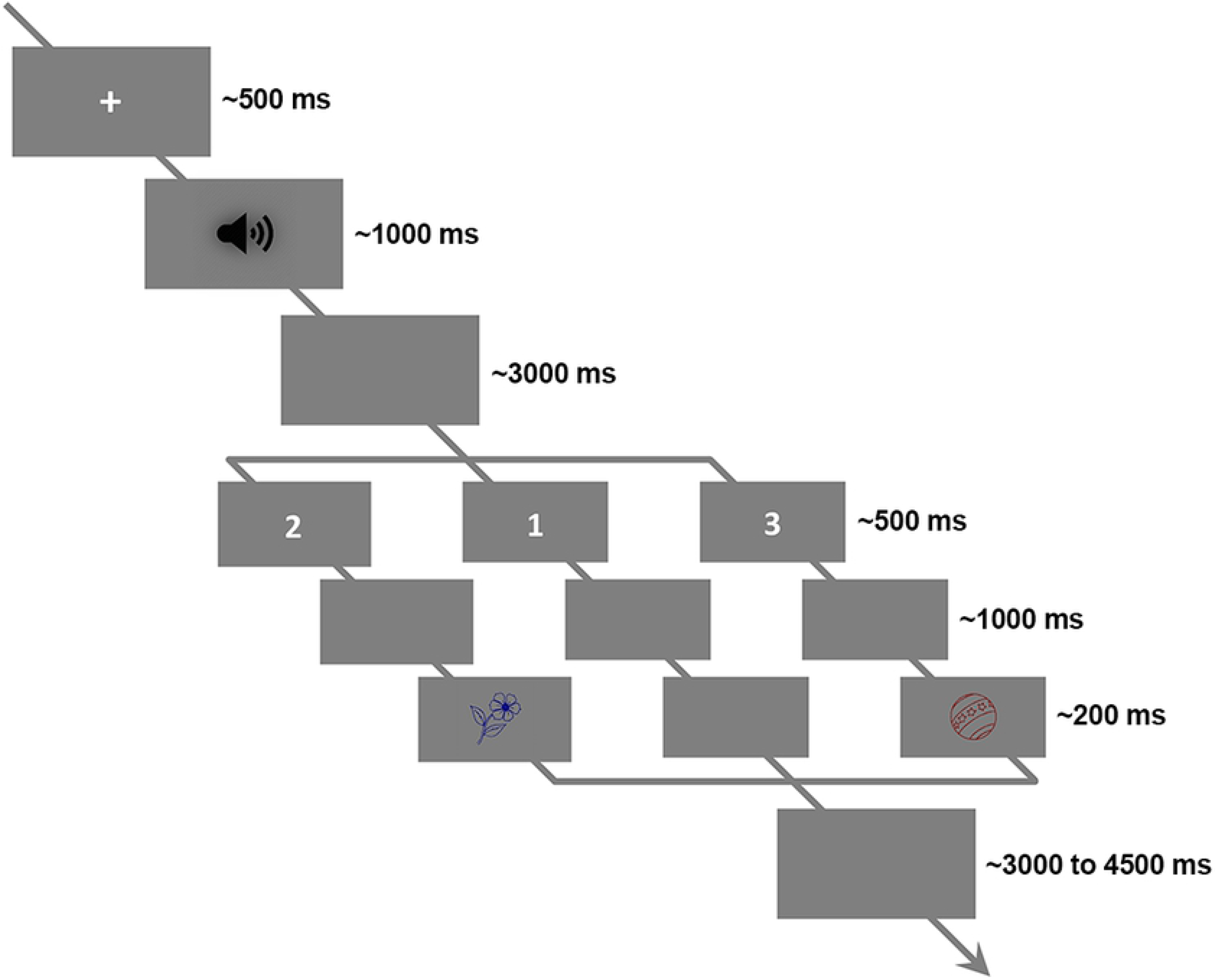
Experimental procedure. An illustration of the experimental procedure. Participants were asked to looking at the screen center, after hearing the speech stimulus they would do a task indexed by a number that showed on the screen. If the number was ‘1’, they would judge whether the heard stimulus was a phrase or a sentence. If the number was ‘2’, they would see a following picture then judge whether the color of the picture was the same as the color that described in the speech stimulus. If the number was ‘3’, they would do an object matching task, in which they would judge whether the object of the picture was the same as the object that described in the speech stimulus. Trial type was pseudo randomly assigned throughout the whole experiment.

Data collection was broken up into 5 blocks, with 48 trials in each block. Before the core data collection, several practice trials were conducted for each participant in order to make sure they had understood the task. Trials in each block were fully matched in across linguistic-structure (phrase or sentence) and task-type (type 1, type 2 or type 3). For instance, half of the spoken stimuli were phrases and half were sentences (twenty-four in each type), six types of combinations (8 trials for each type) were evenly distributed in each block (8*2*3) etc. Trial order was pseudo random throughout the whole experiment. The behavioral results indicated the task was relative easy and no difference between phrase and sentences. All tasks combined accuracy for phrases and sentences were 97.9 ± 3% and 97.3 ± 3% (p = 0.30), respectively.

After the main experiment, a localizer task was performed, in which a tone beep (1 kHz, 50ms in duration) was played 100 times (jitter 2 to 3 seconds) for each participants to localize the canonical auditory response (N1-P2 complex). The topographies for N1 and P2 are shown in **Fig 9**. The upper panel shows the averaged N1-P2 complex of all participants over the time bin from 90 - 110 ms for the N1 and 190 - 210 ms for the P2. The lower panel shows the N1-P2 complex after the Surface Laplacian (Cohen, 2014; Perrin, Bertrand, & Pernier, 1987; Perrin, Pernier, Bertnard, Giard, & Echallier, 1987), in which the effect of the volume conduction was attenuated. The topographies indicated that all participants had the canonical auditory response.

**Fig 9.**
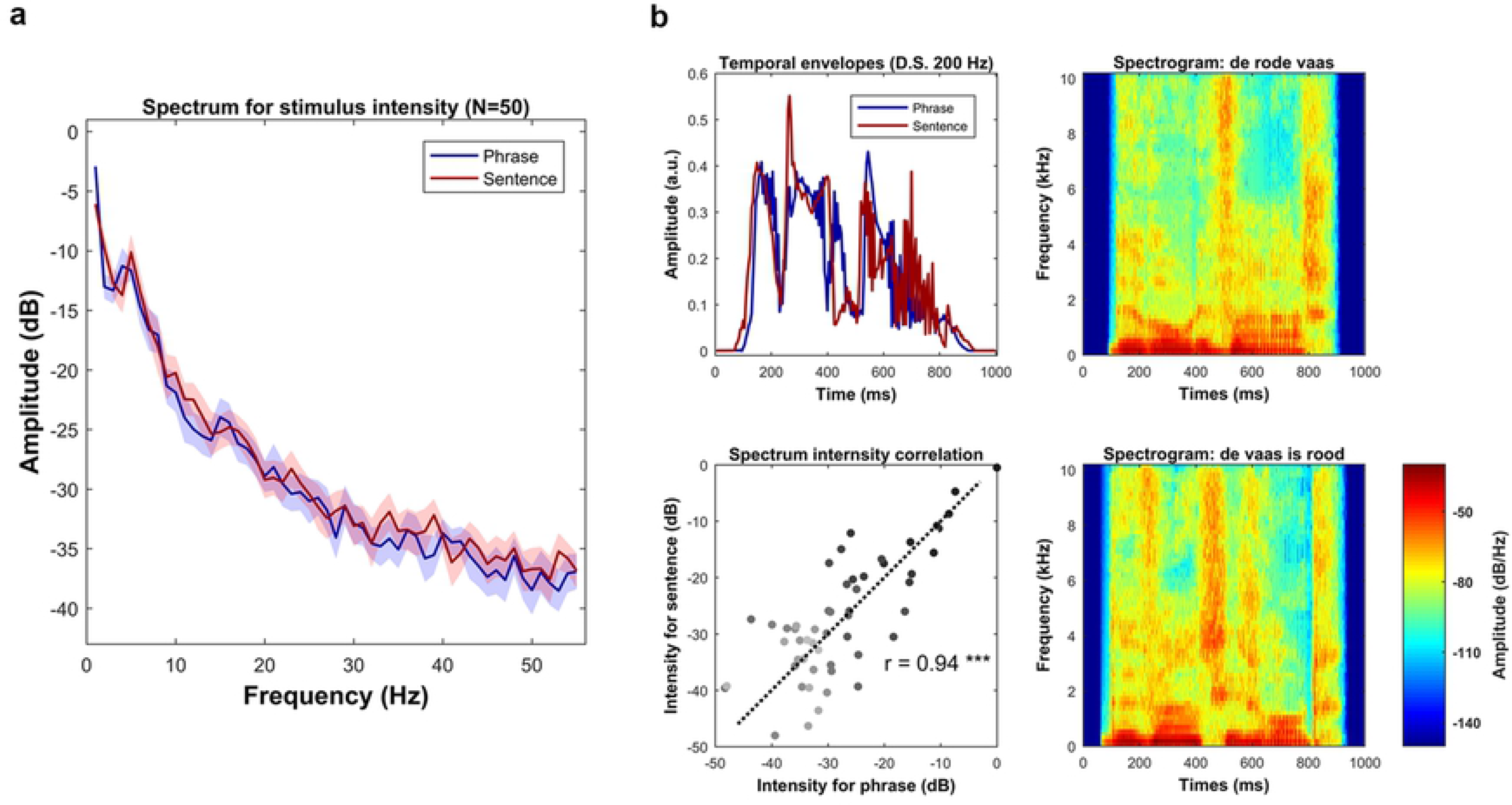
Auditory N1-P2 complex. The topographical distribution of the canonical auditory N1-P2 complex. The upper panel shows the N1-P2 complex that was averaged over all participants (N=15). The lower panel shows the N1-P2 complex after Surface Laplacian (Current Source Density, CSD), in which the effect of volume conduction was attenuated.

### EEG recording

We recorded EEG data using a 64-channel active sensors system of Brain Products (Brain Products GmbH) in a sound dampened, electrically shielded room. Signals were digitized online at 1000 Hz, high-pass and low-pass at 0.01 Hz and 249 Hz, respectively. Two electrodes, AFz and FCz, were used as ground and reference. All electrodes were placed on the scalp based on the international 10-20 system and the impedance of each one was kept below 5 kΩ.

We presented stimuli using MATLAB 2019a (The MathWorks, Natick, MA) with the Psychtoolbox-3 (Brainard, 1997). Auditory stimuli were played at 65 dB SPL and delivered through air tubes ear plugs (Etymotic ER-3C, Etymotic Research, Inc.). Event markers were sent via a parallel port for tagging the onset of the interested events (i.e., speech onset, task index onset, etc.).

### EEG data preprocessing

We preprocessed EEG data via MATLAB using EEGLAB toolbox (Delorme & Makeig, 2004) and customized scripts. We first down-sampled the data to 256Hz, then high-pass filtered at 0.5 Hz (finite impulse response filter, FIR; zero-phase lag), and cleaned by the Artifact Subspace Reconstruction (ASR) (Chang, Hsu, Pion-Tonachini, & Jung, 2018; Kothe & Jung, 2016). We interpolated all detected bad channels with spherical interpolation. After transfer data to average reference, the online reference FCz was recovered and the line noise, 50 Hz and its harmonics, was removed.

Following the above mentioned steps, we extracted epochs of 2 s preceding and 9 s after auditory stimulus onset. We conducted deletion of bad trials deletion and artifacts in the following two steps. First, we used independent component analysis (ICA) for decomposing data to component space (number of components equals data rank). Then for each independent component, we used short-time Fourier Transform (STFT) to convert each trial into power spectrum, in which we extracted a value which was calculated by the power summation between 15 Hz to 50 Hz. Then all the extracted values, one value per epoch, in each component formed a distribution. From this distribution, we transformed all the extracted values to z-score, the epochs with the value outside the range of plus and minus three standard deviation were deleted. Second, ICA was conducted again on the trial-rejected data for eye-related artifacts removal and muscular activities elimination. Artifact components were identified and removed by an automatic classification algorithm (Winkler, Haufe, & Tangermann, 2011). All the preprocessing steps resulted in the removal of, on average, 7 components (range 4 to 11) and 22 trials (including incorrect trials and trials with too slow response, range 10 to 30, 4% to 12.5%) per participant. Finally, volume conduction was attenuated by applying the Surface Laplacian (Cohen, 2014; Perrin et al., 1987; Perrin et al., 1987).

### EEG data analysis

#### Time Frequency Decomposition

We convolved the single-trial time series with a family of complex wavelets (1 to 50 Hz in 70 logarithmically spaced steps), optimizing temporal and spectral resolution by changing the cycle from 3 to 30 in logarithmical steps.

We calculated phase coherence by inter-trial phase clustering (ITPC) ( Cohen, 2014; Lachaux, Rodriguez, Martinerie, & Varela, 1999). At each time-frequency bin, the wavelet coefficients of all trials were divided by their corresponding magnitude and averaged across trials. The magnitude of the averaged complex output was represented as phase coherence (ITPC).

The induced neural response (power) was extracted from the analytical output at each time-frequency bin by taking the summation of the squared wavelet coefficients. Decibel transformation was performed separately at each frequency, in which the average power at the duration from 800 ms to 200 ms before the audio onset was used as the baseline.

#### Phase Connectivity

We first decomposed trials in each condition via wavelet convolution (same parameters as the time-frequency decomposition). Then, we calculated the Cross Spectral Density (CSD) for each sensor-pair at each frequency-time-trial bin. We calculated phase connectivity over the sensor space by Inter Site Phase Coherence (ISPC) (Cohen, 2014; Lachaux et al., 2000; Lachaux et al., 1999; Mormann et al., 2000), in which we divided the complex coefficients by the corresponding amplitude at each frequency-time-trial bin then computed averages across all trials. The amplitude of the averaged complex vector was represented as phase connectivity between sensors (ISPC). After the above mentioned steps, the phase connectivity at each time-frequency bin was represented as an all-sensor to all-sensor matrix, in our case, 65 sensors * 65 sensors.

To transform the connectivity matrix to connectivity degree at each time-frequency bin, a statistical thresholding method was conducted. More specifically, at each time-frequency bin, we formed a distribution by pooling together all the connectivity values from both conditions, then defined the threshold as the value which is half a standard deviation above the median. We then binarized the connectivity matrices for both conditions by applying this threshold at each bin. The connectivity degree at each time-frequency bin was then represented as the count of the number of the connectivity values that above this threshold. Finally, we normalized the connectivity level at each time-frequency bin to percentage change relative to the baseline which was calculated as the average connectivity degree at the duration from 800 ms to 200 ms before the audio onset.

#### Phase Amplitude Coupling (PAC)

Since low-frequency phase and high-frequency amplitude may show coupling during speech processing (Giraud & Poeppel, 2012), we defined the frequency range for the phase time series from 1-16 Hz in a linear step of 1.5 Hz, and the frequency range for the amplitude time series from 8 Hz to 50 Hz in 12 logarithmically steps. Then, we performed the wavelet convolution to extract the analytic signals, in which we extracted the phase time series and amplitude time series at the specified frequency range from 50 ms before to 1500 ms after the audio onset. At each phase-amplitude bin, we constructed a complex time series that held the phase angle of the phase time series and weighted by the magnitude of the amplitude time series. We calculated the PAC at each bin by extracting the magnitude of the average of all the vectors in the complex time series (Canolty et al., 2006; Cohen, 2014). Since the variation of the amplitude response, a Z-score normalization was also performed for each phase-amplitude bin. More specifically, after calculating the real PAC value using the raw complex time series, the random PAC value were computed 1000 times using the re-constructed complex time series, which were built by temporally shifting the amplitude time series with a random temporal offset. These 1000 random PAC values formed a reference distribution for each phase-amplitude bin. Then the z-score of the real PAC value in this distribution was represented as the index of the phase-amplitude coupling, PAC-Z.

#### Power Connectivity

After time-frequency decomposition, we extracted induced power at each channel-time-frequency-trial bin. For each condition, we calculated power connectivity for each sensor-pair at each time-frequency bin as the Rank Correlation between the power response of all trials in one sensor and the power response of all trials in the other sensor (Bruns, Eckhorn, Jokeit, & Ebner, 2000; Cohen, 2014; Hipp, Hawellek, Corbetta, Siegel, & Engel, 2012). The power connectivity calculation resulted in an all-sensors to all-sensors (65*65 in our case) representation at each time-frequency bin for each condition.

To transfer the power connectivity at each time-frequency bin to the power connectivity degree, we applied a statistical thresholding method. More specifically, at each time-frequency bin, we formed a distribution by pooling together all the connectivity values from both conditions, then defined the threshold as the value which is half a standard deviation above the median. We then binarized the connectivity matrix at each bin for each condition by applying the corresponding threshold. The connectivity degree at each time-frequency bin was represented as the count of the number of the connectivity values that above this threshold. Finally, we transferred the connectivity level at each time-frequency bin as a percentage change relative to the connectivity level of the baseline which was calculated as the average connectivity level in the duration from 800 ms to 200 ms before the audio onset.

#### Spectral Temporal Response Function (STRF)

The STRF is a linear kernel which convolves with the specified features of the speech signal to estimate the neural response in time. It can be interpreted as a linear filter which transforms the stimulus feature to the neural response (Crosse, Di Liberto, Bednar, & Lalor, 2016; Di Liberto et al., 2015).

In our study, the stimulus feature (SF) was defined as the spectrogram, which was obtained by filtering the speech stimulus into 16 logarithmically spaced frequency bands between 0.05 to 8 kHz for mimicking the frequency decomposition by the brain (Greenwood, 1990). The temporal envelope (energy profile) for each frequency band was then extracted by Hilbert Transform.

To construct the stimulus-response pairs, we first applied a linear ramp to both sides of each trial corresponded neural response (10% at each side) to attenuate the abrupt onset and offset. Then, we matched each 1-second neural response with the corresponding SF.

In order to optimize the estimation of the STRF, a randomization procedure was applied to create new data structure. We first randomly selected 80% of all unique speech stimuli, then the stimulus-response pairs that corresponded to the selected speech stimuli were extracted as the seed data to construct the training dataset for performing the cross validation. We constructed thirty five 10-second long stimulus-response pairs, in which each one of them was a concatenation of the randomly selected ten 1-second stimulus-response pair (Bootstrapping).

The STRF was estimated using the Ridge Regression with leave-one-out cross validation. Since the Ridge Regression weights the diagonal elements of the covariance matrix of the SF with a parameter Lambda (Crosse et al., 2016; Tikhonov & Arsenin, 1977), we, therefore, predefined the range of Lambda to 10 values from 6 to 100 in linear steps. We used the extracted training dataset (thirty five 10-second long stimulus-response pairs) to conduct the cross validation for optimizing the STRF. The Pearson’s correlation between each real neural response and each predicted response was calculated. The average of all the coefficients of the Pearson’s correlation (over all sensors and all trials) was defined as the performance of the STRF. The model with the Lambda parameter which gave the best performance was used as the optimized STRF.

Since previous research has shown that slow timescale (low frequency) neural response reliably reflects the neural representation of the acoustic features in speech (Ding & Simon, 2012a, 2012b), we, therefore, initially checked whether the STRF for different frequency bands faithfully reflect the encoding of the acoustic features. To do so, we first filtered the neural response corresponding to each trial into five canonical frequency bands, which were Delta (< 4 Hz), Theta (4 to 7 Hz), Alpha (8 to 13 Hz), Beta (14 to 30 Hz) and Low-Gamma (31 to 50 Hz), respectively. Then, the STRF for each condition at each frequency band was estimated using the procedure that mentioned above. In order to check the performance of each estimated STRF, the stimulus-response pairs (the unseen pairs for each STRF) that corresponded to the remaining 20% of the speech stimuli were extracted as the seed data for constructing the testing dataset. We extracted five 4-second long stimulus-response pairs in the testing data for each STRF, each one of them was a concatenation of randomly selected four 1-second long pairs.

The real performance of each STRF was calculated by using the frequency and condition matched stimulus-response pairs. The random performance was calculated 1000 times using the pairs which were constructed by combining the stimulus with a randomly selected neural response.

For fitting the low frequency STRF (< 9 Hz), the same procedure was conducted. The performance of the averaged STRF in each condition was computed using the averaged Pearson’s correlation between the real neural response and the predicted response across sensors.

We extracted the temporal response function (TRF) and the spectral response function (SRF) for each participant at each condition by averaging the STRF over the frequencies from 0.1 kHz to 800 kHz and by averaging the STRF over the time from 0 to 400 ms, respectively.

All the calculations in this section were conducted using customized scripts, the scripts of EEGLAB toolbox (Delorme & Makeig, 2004), and the Multivariate Temporal Response Function Toolbox (Crosse et al., 2016).

#### Statistical Analysis

In addition to using parametric statistical methods, we applied a cluster-based nonparametric permutation test. This method deals with the multiple-comparisons problem and at the same time takes into account the dependencies (temporal, spatial and spectral adjacency) in the data. For all types of analysis that followed this inference method, we initially averaged the subject-level data over trials and for each single sample, i.e. a time-frequency-channel point, and performed a dependent t-test. We selected all samples for which the t-value exceeded an a priori threshold, p<0.05, two-sided, and subsequently clustered these on the basis of spatial and temporal-spectral adjacency. The sum of the t-values within a cluster was used as cluster-level statistic. The cluster with the maximum sum was subsequently used as the test statistic. By randomizing the data across the two conditions and recalculating the test statistic 1000 times, we obtained a reference distribution of the maximum cluster t values. This distribution was used to evaluate the statistical distribution of the actual data. This statistical method was implemented using the FieldTrip toolbox (Maris & Oostenveld, 2007; Oostenveld, Fries, Maris, & Schoffelen, 2011).

#### Acoustic analysis

The intensity fluctuation of each speech stimulus was characterized by the corresponding temporal envelope, which was extracted by the Hilbert Transform of the half-wave rectified speech signal. Then each extracted temporal envelope was downsampled to 200 Hz and the Discrete Fourier Transform (DFT) was performed to extract the spectrum. Decibel transformation for the spectrum of each speech stimulus was performed by using the highest frequency response in the corresponding phrase-sentence pair as the reference. **Fig 10a** shows the comparison between the averaged spectrum of all phrases and the averaged spectrum of all sentences. The shaded area covers 2 standed errors of the mean across the stimuli. Statistical comparison (the paired-sample t-test) was conducted at each frequency bin, in which no evidence was found to indicate significant physical difference between phrases and sentences. **Fig 10b** shows the acoustic features of a phrase-sentence pair, *De rode vaas (The red vase) and De vaas is rood (The vase is red),* the upper-left panel shows the comparison of the temporal envelopes of this sample pair. The lower-left panel shows the intensity relationship of this sample pair in each frequency bin. The Pearson’s correlation was calculated to reveal the similarity between the spectrum of this sample pair (r = 0.94, p < 1e-4). The comparisons indicated that they are highly similar in acoustic features. In this figure, the darker the dots represents the lower the frequency. The upper-right panel and lower-right panel shows the spectrogram the sample phrase and the sample sentence, respectivily.

**Fig 10.**
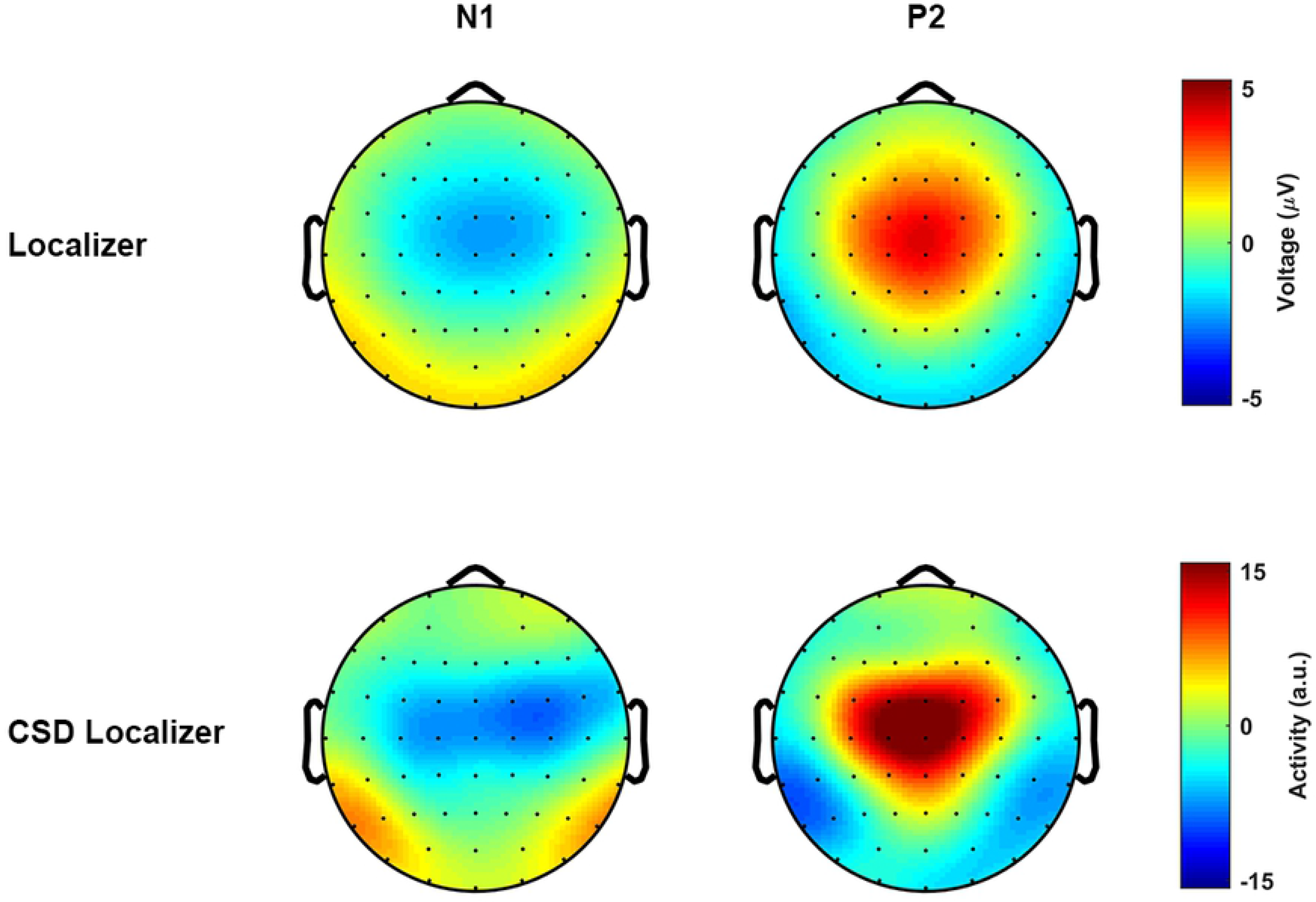
Stimuli comparison. **(a)** Spectrum comparison between the two types of speech stimuli. The shaded area for each condition covers 2 s.e.m. across the mean (N=50). **(b)** The upper-left panel, shows the comparison of the temporal envelopes of the sample phrase-sentence pair, i.e., *De rode vass (the red vase)* v.s. *De vaas is rood (the vase is red)*, that down sampled to 200 Hz. The lower-left panel shows the spectrum for the sample phrase-sentence pair, in which the horizontal axis and the vertical axis shows the frequency response of the temporal envelop of the phrase and the sentence, respectively. The Pearson’s correlation indicated that the spectrum are highly similar between the sample phrase and the sample sentence (r = 0.94, p < 1e-5 ***). The upper-right and lower-right panel shows the spectrogram of the sample phrase and the sample sentence, respectively.

## Discussion

In this study, we investigated the neural responses to minimally different linguistic structures by minimizing their differences in acoustic-energetic/temporal-spectral profiles and semantic components. We investigated which dimensions of neural activity distinguish the linguistic structure of phrases and sentences and used a series of analysis techniques to better describe the dimesion of neural readouts that were sensitive to the distinctive linguistic structures between phrases and sentences. We asked whether phrases and sentences have different effects on functional connectivity, and found, first, that while phrases and sentences recruit similar functional networks, the engagement of those networks scaled with linguistic structure. Sentences showed more phase coherence and power connectivity compared to phrases. This connectivity pattern suggests that phrases and sentences differently impact the distribution and intensity of neural networks involved in speech comprehension. Second, we found that phase-amplitude coupling between theta and gamma, which has been implicated in speech processing, is not sensitive to structural differences in spoken language. Third, we found that activity in the alpha band was sensitive to linguistic structure. Lastly, by modelling acoustic fluctuations in the stimulus and brain response with STRFs, we found that phrases and sentences differentially relied on the acoustic response in the brain, and that sentences were more abstracted away from acoustic dynamics in the brain response. In the following sections we give more detail about our findings and discuss potential interpretations of them.

### Phase coherence

Consistent with previous research (Doelling et al., 2014; Luo & Poeppel, 2007; Peelle & Davis, 2012; Peelle et al., 2013), our phase synchronization analysis detected low frequency phase coherence during speech comprehension. Moreover, phase coherence distinguished between phrases and sentences, yielding a cluster between ∼450 and ∼900 ms after audio onset and from ∼2 Hz to ∼8 Hz in frequency, that was most pronounced over central electrodes. These results therefore suggest that syntactic structure may be encoded by low frequency phase coherence, through systematic organization of activity in neural networks, in particular their temporal dynamics. Our results are consistent with the notion of *phase sets* in computational models of structured representations that exploit oscillatory dynamics. Phase sets are representational groupings that are formed by treating distributed patterns of activation as a set when units are in (or out) of phase with one another across the network (Doumas & Martin, 2018; Martin & Doumas, 2017, 2019, 2020). They are key to the representation of structure in artificial neural network models.

### Phase connectivity

Phrases and sentences also yielded differences in phase connectivity. In the predefined time and frequency range of interest, the statistical comparison indicated a difference corresponding to a cluster approximatly from ∼800 to ∼1600 ms after the audio offset, occurring at the very low frequency range (< ∼2 Hz) that was most pronounced over the right posterior region. Phrases and sentences thus differentially impact the temporal synchronization of neural responses.

Several aspects of the results are noteworthy. First of all, the relatively late effect suggests that the effect on temporal synchronization occurs after the initial presentation the speech stimulus. In our experiment, participants were randomly presented a task prompt for three possible tasks (color discrimination, object discrimination, phrase or sentence discrimination), which asked them to determine either ‘semantic’ (object or color information) or ‘syntactic’ information (whether the stimulus was a phrase or sentence) from the speech stimulus. Because of the random order of the task trials, participants had to pay close attention the stimuli and continue to represent each stimulus after hearing it, namely until they received the task prompt. The tasks also insured that participants could not select a single dimension of the stimulus for processing. Similarly, because we used an object and a color task, participants had to distribute their attention evenly across the adjectives and nouns, mitigating word order differences between structures. In light of these controls and task demands, we consider it unlikely that the observed phase connectivity effects reflect mere differences in attention to phrases or sentences. Rather, we attribute the observed effects to the syntactic differences between them.

Secondly, the low frequency range (< 2 Hz) of the observed effect is consistent with previous research (Brennan & Martin, 2020; Ding et al., 2016; Kaufeld et al., 2020; Keitel et al., 2018; Meyer et al., 2017). In Ding et al. (2016), the cortical response was modulated by the timing of the occurrence of linguistic structure; low frequency neural responses (1-2 Hz) were found to track the highest level linguistic structures (phrases and sentences) in their stimuli. Here we extended their work to ask whether the 1 Hz response could be decomposed to reflect separate syntactic structures (phrases vs. sentences), and we identified the role of phase in discriminating between these structures. In our study, all speech stimuli lasted 1 second, and except for the presence of syntactic structure, the stimuli were normalized to be highly similar. Our pattern of results therefore suggests that functional connectivity, as reflected in the temporal synchronization of the induced neural response, distinguishes between phrases and sentences.

Lastly, phrases and sentences differed most strongly over right-posterior parietal cortex, which is broadly consistent with previous research on speech comprehension. Functional Magnetic Resonance Imaging (fMRI) studies implicate the posterior right hemisphere in processing syntactic structure (de Bode, Smets, Mathern, & Dubinsky, 2015; Grodzinsky, 2000; Grodzinsky & Friederici, 2006; Maess, Koelsch, Gunter, & Friederici, 2001). Neurophysiological research also suggests the involvement of right hemisphere in extraction of slow timescale information extraction (Abrams, Nicol, Zecker, & Kraus, 2008; Giraud et al., 2007; Morillon, Liégeois-Chauvel, Arnal, Bénar, & Giraud, 2012; Poeppel, 2003). In addition, the P600, a positive ERP component often associated with syntactic processing, has a robust right posterior topographical distribution (Coulson, King, & Kutas, 1998; Friederici, Pfeifer, & Hahne, 1993; Hagoort, Brown, & Groothusen, 1993; Osterhout & Holcomb, 1992; Osterhout & Mobley, 1995; Patel, Gibson, Ratner, Besson, & Holcomb, 1998). In light of the existing literature, therefore, the right-posterior distribution of the phase connectivity effects is consistent with processing of syntactic structures, although we refrain from claims about underlying neural sources based on our EEG data.

### Phase Amplitude Coupling (PAC)

We observed PAC during speech comprehension, as a low frequency phase response (∼ 4 – 10 Hz) entrained with high frequency amplitude (∼ 15 – 40 Hz). This effect appeared largely over the bilateral central area. The bilaterial central topographical distribution has been replicated shown reflecting sensory-motor integration (Babiloni et al., 2011; Klimesch, Sauseng, & Hanslmayr, 2007; Neuper, Wörtz, & Pfurtscheller, 2006; Pfurtscheller, Brunner, Schlögl, & Da Silva, 2006; Pfurtscheller, Neuper, Schlogl, & Lugger, 1998; Schlögl, Lee, Bischof, & Pfurtscheller, 2005; Suffczynski, Kalitzin, Pfurtscheller, & Da Silva, 2001), which is consistent with the proposal from Giraud and Poeppel (2012) that PAC reflects an early step in speech encoding involving sensory-motor alignment between the auditory and articulatory systems. Crucially, however, this effect did not distinguish phrases and sentences. Although a null result, this pattern is compatible with the generalized model for speech perception (Giraud & Poeppel, 2012). This early step is presumably similar for phrases and sentences, and perhaps for any type of structure above the syllable level.

### Induced alpha power

Induced alpha band power distinguished phrases and sentences, and this effect was most pronounced at the left hemisphere. This pattern suggests involvement of alpha band oscillations in syntactic structure processing. Although alpha band activity is often associated with attentional or working memory related processing (Haegens et al., 2010; Obleser et al., 2012; Strauß et al., 2014; Ten Oever et al., 2020; Wilsch & Obleser, 2016; Wöstmann et al., 2016; Wöstmann et al., 2015; Wöstmann et al., 2017), we do not consider this a very plausible alternative explanation for our results. For example, it not clear why phrases and sentences would differ in their attentional demands. In addition, we employed an experimental task to ensure similar attention to phrases and sentences, and phrases and sentences were associated with similar behavioral performance in each task (with a caveat that performance was at ceiling and may therefore not pick up on small differences between conditions).

We do not claim that that all speech-elicited alpha band effects reflect syntactic processing. Some observed effects effects may well reflect perceptual processing during speech comprehension (e.g. Obleser & Weisz, 2012), especially in experiments designed to manipulate perceptual processing, such as speech-in-noise manipulations. Neural response in a given band, e.g., the alpha band, need not reflect only a one particular perceptual process. Likewise, the fact that the alpha band neural response could reflect lower-level perceptual processes or working memory load in certain contexts does not necessarily rule out its role in higher-level linguistic information representation, e.g., syntactic information.

### Power connectivity

Phrases and sentences elicit differences in induced power connectivity in alpha band activity (7.5 to 13.5 Hz). Phrases showed more inhibition in power connectivity than sentences; in other words, sentences showed a stronger connectivity degree over sensor space in the alpha band than phrases. Several aspects of these results are noteworthy. First, we observed this effect from ∼100 ms until ∼2200 ms after stimulus onset, which suggests the effect in functional connectivity persisted rather long and outlasted the observed effects in induced alpha power, which we observed from ∼350 ms to ∼1000 ms after the audio onset (during the listening stage). The extended nature of the functional connectivity effect could reflect the continuing integration and representation of syntactic and semantic components.

Secondly, alongside differences between phrases and sentences in power connectivity, we also extracted the sensor connectivity pattern (over an ROI ranging from 100 ms to 2200 ms in time and 7.5 to 13.5 Hz in frequency). Whereas phrase and sentences showed similar functional connectivity in the intensity of the neural response, sentences showed stronger inter-region (sensor) connectivity than phrases. By design, in our stimuli sentences had more constituents than phrases did. If local network activity is more organized or coherent as a function of linguistic structure, then the difference observed here could reflect the encoding of additional constituents in sentences compared to phrases.

Lastly, phrases elicit stronger inhibition of induced power connectivity than sentences did. This indicates weaker cooperation between brain regions, in other words, regions showing more independent the neural response. In contrast, inter-region connectivity was stronger for the sentence condition than the phrase condition, which suggested a higher-level of intensity of connectivity between brain regions for the sentences in order to separate them from the phrases.

In sum, phrase and sentences elicited robust differences in induced power connectivity. A similar functional connectivity pattern was deployed for representing phrases and sentences, but the intensity of the connectivity was stronger for sentences than phrases. This finding is consistent with the prediction that low frequency power and network ogranization should increase as linguistic structure increases. Our stimuli were designed to allow the measurement of differences in neural dynamics between phrases and sentences, and as such differed in the number and type of linguistic constituents that were perceived. Beyond the number and type of consituents, the phrase and sentence structures also differ in the relations between constiuents, or in the linguistic notion of hierarchy. But given the co-extension of number, type, and relation, our stimuli and design do not yet allow us to determine if, nor how, these variables alone might affect structural encoding in neural dynamics.

### Spectrotemporal response functions (STRF)

We performed STRF analysis to investigate whether phrases and sentences are encoded differently. Firstly, consistent with previous research (Ding & Simon, 2012a, 2012b), only low frequency (< 9 Hz) neural responses robustly reflected the phase-locked encoding of the acoustic features. Secondly, we observed a bilaterial representation of the slow temporal modulations of speech, in particular at posterior sensors. The posterior region has consistently been found to be involved in syntactic integration (Coulson et al., 1998; Friederici et al., 1993; Hagoort et al., 1993; Osterhout & Holcomb, 1992; Osterhout & Mobley, 1995; Patel et al., 1998). The low frequency neural response that models the phase-locked encoding of acoustic features can capture structural differences between phrases and sentences, even without explicitly using a hand-coded annotation on the syntactic level to reconstruct the data. Thirdly, and most importantly, we explored these patterns further in both the temporal and spectral dimension, by decomposing the STRF into the TRF and the SRF. Both TRF and SRF suggested a different encoding mechanism between phrases and sentences. More specifically, the TRF results showed that the brain transduces the speech stimulus into the low frequency neural response via an encoding mechanism with two peaks in time (at ∼100 ms and ∼300 ms). The two peaks reflect the instantaneous low frequency neural response that is predominantly driven by the encoding of acoustic features that were presented ∼100 ms and ∼300 ms ago. In the two time windows that centered at ∼100 ms and ∼300 ms, respectively, phrases and sentences showed a different dependency on the acoustic features in both latency and intensity.

When we only consider intensity (∼100 ms time window), sentences depended on acoustic features less strongly than phrases. This result is consistent with the idea that sentence representations are more abstracted away from the physical input because they contain more linguistic structural units (i.e., constituents) that are not verdically present in the physical or sensory stimulus. Consistent with previous research, we found that the instantaneous neural response was strongly driven by the encoding of the acoustic features presented ∼100 ms ago (Brodbeck et al., 2018; Crosse & Lalor, 2014; Di Liberto et al., 2015; Ding & Simon, 2012a, 2012b, 2013; Golumbic et al., 2013; Puvvada & Simon, 2017; Wang, Zhang, Zou, Luo, & Ding, 2019).

When we only consider the latency (∼100 ms time window), and only at the right hemisphere, the low frequency neural response of the sentences was predominantly driven by the acoustic features that appeared earlier in time than the acoustic features that drove the neural response to phrases. Our results imply that the brain distinguishes syntactically different linguistic structures by how driven its responses are by the acoustic features that appeared at ∼100 ms ago. More importantly, at the right hemisphere, the findings suggested that the low frequency neural response of the sentences reflected the encoding of the acoustic features that appeared earlier in time than the acoustic features that drove the neural response of the phrases. This could be evidence that the right hemisphere is dominant in extracting the slow timescale information of speech that is relevant for, or even shapes, higher-level linguistic structure processing, e.g., syntactic structure building (Ding & Simon, 2012a, 2012b; Poeppel, 2003). It is noteworthy to see that the distribution in time and space of these patterns is consistent with the idea that the brain is extracting information from the sensory input at different timescales, and that this process is in turn is reflected in the degree of departure (in terms of informational similarity) of the neural response from physical features of the sensory input.

At ∼300 ms, when we only consider intensity, the low frequency neural response to phrases is more strongly dependent on the acoustic features than the neural response to sentences is, showing again that sentences are more abstracted from sensory representations. However, the comparison of the TRF indicated that the brain exploited a different encoding mechanism at the right hemisphere between the phrases and the sentences. More concretely, the low frequency neural response of the phrases showed a stronger dependency on the acoustic features than the sentences at the right hemisphere, but not at the left hemisphere. The results, first, suggested that the instantaneous low frequency neural response reflects the encoding of the acoustic features that were present ∼300 ms ago, in both conditions. Then, it reflected that, only in the right hemisphere, the low frequency neural response of the phrases more strongly depends on the acoustic features from ∼300 ms ago when compared to the neural response to sentences. This finding reminds us of the results of the phase connectivity analysis, in which the phase connectivity degree also showed a different pattern between the phrases and the sentences at the right posterior region; which is consistent with previous research that has shown a role for the right hemisphere in processing slow modulations in speech, such as prosody (Abrams et al., 2008; Ding & Simon, 2012a, 2012b; Giraud et al., 2007; Kerlin, Shahin, & Miller, 2010; Luo & Poeppel, 2007; Poeppel, 2003). Consistent with these findings, our results further indicated that the brain can separate syntactically different linguistic structures via differential reliance by the right hemisphere on representations of the acoustic features that appeared at ∼300 ms ago. That sentence representations were more abstract and less driven by the acoustics in the left hemisphere is consistent with contemporary neurobiological models of sentence processing (Friederici, 1995; Hagoort, 2013), although we cannot attribute the differences we observed to a single aspect of linguistic structural descriptions (i.e., to number, type, relations between constituents, or whether constituents form a ‘saturated’ sentence or not).

The SRF results indicated that the brain can begin to separate phrases and sentences via differential reliance on the encoding of the acoustic features from roughly the first formant (Catford, 1988; Jeans, 1968; Titze et al., 2015; Titze & Martin, 1998), and in a phase-locked manner. More specifically, in the range of the first formant, the low frequency neural response reflected a stronger dependency on the acoustic features in the phrases than in the sentences. Unlike consonants, the intensity of vowels are well reflected at the first formant (<1 kHz) (Catford, 1988; Jeans, 1968; Titze et al., 2015; Titze & Martin, 1998). Although the overall physical intensity of the speech stimulus of the phrases was not different from the sentences, the neural response contains information that discriminates the differences between syntactic structures. Given that the stimuli were not physically different, this pattern of results strongly suggests that the brain is “adding” information, for example by actively selecting and representing linguistic structures that are cued by the physical input and its sensory correlate. For example, the brain could be adding phonemic level information, e.g., vowels, lexical, and phrasal via a top-down mechanism; in certain situations and languages, even a single vowel can cue differential syntactic structure. In fact, in our stimuli, the schwa carries agreement information that indicates the phrasal relationship between “red”(*roode)* and “vase”(vaas) in the phrase “de roode vaas.” Our results, which feature both dependence on, but also departure from, the acoustic signal, are consistent with previous findings that have shown low frequency cortical entrainment to speech and argued that is can reflect phoneme-level processing (Di Liberto & Lalor, 2017; Di Liberto et al., 2015; Keitel et al., 2018; Khalighinejad, da Silva, & Mesgarani, 2017). We extend these findings by showing that when lower-level variables in the stimuli are modeled, the brain response can discriminate between syntactic structures even without the addition of higher-level linguistic annotations in our TRF models, and that this result indicates that the degree of departure from the physical stimulus increased as abstract structure accrues.

### Summary

In this study, we found a neural differentiation between spoken phrases and sentences that were physically and semantically similar. Moreover, we found that this differentiation was captured in several readouts, or dimensions of brain activity (*viz.*, phase synchroniziation, functional connectivity in phase and induced power). By modeling the phase-locked encoding of the acoustic features, we further show that the brain can represent the syntactic difference between phrases and sentences in the low frequency neural response, but that the more structured a stimulus is, the more it departs from the acoustically-driven neural response to the stimulus, even when the physicality of the stimulus is held constant. Across all our results, we provide a comprehensive electroencephalographic picture of how the brain separates linguistic structures within its representational repetoire. However, further research is needed to explore the relationship between these different neural readouts that indexed syntactic differences, e.g., how the induced neural response at alpha band interacts with phase coherence in the low frequency (< 8 Hz), and how these separation effects are represented at the neural source level.

## Acknowledgments

AEM was supported by the Netherlands Organization for Scientific Research (NWO; grant 016.Vidi.188.029), and a Max Planck Research Group and a Lise Meitner Research Group “Language and Computation in Neural Systems” from the Max Planck Society. The authors thank Mante S. Nieuwland for comments on an earlier version of this work, Shunag Bi for selecting stimuli and arranging figures, and Cas W. Coopmans for guidance as to visual representation of linguistic constituents.

## Footnotes

1 Their model exploited asynchrony of unit firing to form structures from words, in which the temporal proximity of unit firing was used to encode linguistic structures and relations. More specifically, nodes on higher layer (i.e., localist units representing phrasal units) code for the composition of sub-layer units (i.e., words) in this network structure. The higher-level node fires when any of its sublayer nodes fire in time, forming a ‘phase set’ between words and their phrase. In this implementation, linguistic structures were represented neural dynamics across a layered network, where the asynchrony of unit firing (firing staggered in time) not only allowed the network to combine lower-level representations together for the processing on a higher-level, but also served to keep the lower-level representations independent as well (Doumas et al., 2008; Martin & Doumas, 2017, 2019).

